# A direct forcing immersed boundary method for biofluid simulations using a non-linear rotation free shell model on unstructured grids

**DOI:** 10.64898/2026.05.16.725689

**Authors:** Taeouk Kim, Abhilash R. Malipeddi, Jesse Capecelatro, C. Alberto Figueroa

## Abstract

Thin structures such as heart valves and aortic dissection flaps interact dynamically with blood flow in human vessels. Their flexibility and capacity for large deformations generate complex, highly transient hemodynamic patterns over the cardiac cycle. Accurately resolving these interactions remains challenging for conventional boundary-fitted fluid–structure interaction approaches. We present an immersed boundary method for simulating thin structures in incompressible flow on unstructured grids. The method couples a stabilized finite element fluid solver with a nonlinear, rotation-free shell formulation through a direct forcing immersed boundary approach. The framework supports both weak (explicit) and strong (implicit) time-coupling strategies, enabling stable simulations over a wide range of solid-to-fluid density ratios. Hydrodynamic forces acting on thin structures are computed from fluid solutions sampled on both sides of the structure, allowing accurate force reconstruction for zero-thickness shells. To our knowledge, this is the first immersed boundary formulation that couples an unstructured finite element fluid solver with a two-dimensional, rotation-free shell model to simulate interactions between thin structures and incompressible flow. Fluid–structure coupling is achieved using predefined finite element shape functions, which provide consistent projection between Eulerian and Lagrangian fields without additional interpolation procedures. The framework is validated using three-dimensional benchmark problems involving thin structures. Then, valve-like model is used to compare strong and weak coupling strategies. Finally, the method is applied to an idealized type-B aortic dissection model. The proposed approach is implemented within the open-source software CRIMSON, a finite element platform for cardiovascular simulation.

## 1. Introduction

In the cardiovascular system, interactions between thin structures and blood flow occur both during normal (e.g., heart and venous valves) as well as pathological (e.g., aortic dissection flaps) conditions. These thin structures are flexible and undergo large deformations, resulting in complex blood flow patterns during the cardiac cycle. Understanding the fluid-structure interactions associated with these thin structures is highly relevant in diseases such as aortic insufficiency, venous insufficiency, and dynamic obstruction in aortic dissection [1, 2, 3]. The motion of these structures has been studied using various dynamic medical image techniques, such as computed tomography (CT), magnetic resonance imaging (MRI), and ultrasound [4, 5, 6]. These imaging methods are valuable for disease diagnosis and classification [7, 8, 9, 10]. However, relying solely on medical imaging makes it challenging to gain insights into how these large motions affect physiological function and contribute to disease, because blood flow and pressure data under such conditions are limited due to the constraints of clinical hemodynamic measurements [11, 12]. To address these challenges, computational simulations have been used to investigate the complex biomechanics of thin structures and their impact on blood flow in vessels.

Methods to simulate fluid–structure interaction (FSI) have been used in simulations of the aortic valve and aortic dissection to study the effects of structural motion on hemodynamic parameters. FSI methods can be categorized according to two main criteria: time coupling strategies and interface handling techniques [13, 14]. Regarding time coupling strategies, FSI methods are classified into monolithic or partitioned approaches [14]. In monolithic approaches, fluid and structural equations are solved simultaneously as a single system. This ensures that the resulting fluid-solid interactions are synchronized in time and unconditionally stable [15, 16]. However, monolithic approaches require an integrated system with stabilization terms [17]. In contrast, partitioned approaches solve fluid and structural domains sepa-rately using distinct solvers. This separation introduces a time lag between fluid and solid solutions [14, 18]. To address this issue, sub-iterations between fluid and structural solutions are performed until convergence is achieved for each time step [19, 20]. Partitioned approaches offer greater flexibility, allowing the use of pre-existing solvers for fluid and structural domains with minor modifications [21].

Regarding interface handling techniques, FSI methods differ in the handling of the interface between fluid and structural domains and are classified into boundary-fitted (conforming mesh) and non-boundary-fitted (non-conforming mesh) methods. In boundary-fitted methods, the fluid is solved on a moving mesh that follows the motion of the solid. The Arbitrary Lagrangian–Eulerian (ALE) method is a prominent example [22]. ALE allows for strict imposition of boundary conditions at the fluid-solid interface. However, ALE methods require additional computational cost for re-meshing, possibly at each time step, and may suffer from mesh distortion under large structural motions within short time frames [23, 24]. ALE methods have been used for simulations of deformable aortic walls [25, 26, 27] and aortic dissection [28, 29, 30]. In contrast, non-boundary-fitted methods use a fixed Eulerian grid for the fluid domain while representing the solid domain using a separate Lagrangian solver. Immersed boundary methods (IBM), originally introduced by Peskin [31], are a family of non–boundary–fitted formulations widely used in FSI. These methods avoid re-meshing and are well suited for handling complex structural geometries and large deformations [31, 32] using dedicated solid solvers. Boundary conditions at the interface are enforced through body forces in the Navier–Stokes equations near the solid domain. Since the solid boundary does not conform to the fluid grid, IBM methods require interpolation or smoothing to integrate boundary forces with the fluid solver. Various approaches have been developed that differ in body force formulation and interpolation strategy, including continuous forcing [31, 33], sharp-interface [34], and direct forcing approaches [35, 36]. In the context of biofluid simulations, IBM have been successfully applied to aortic valve simulations [37, 38, 39].

Thin structures within blood vessels undergo large deformations over relatively short time scales, such as during the cardiac cycle (*∼*1 second). These structures may also move close to, or even come into contact with, other solid structures, such as aortic valve leaflets during diastole [40] or the intimal flap and the aortic wall in dynamic obstruction [30]. ALE methods are ill-suited to these scenarios because frequent remeshing is computationally expensive and can lead to mesh distortion and degraded mesh quality [41]. In such cases, IBM offers an attractive alternative, as it avoids the need for fluid re-meshing and effectively captures the dynamics interactions between fluid and thin structures.

FSI simulations of biological tissues require the use of robust and efficient structural models that accurately represent structural deformations. While membrane models are a popular choice for modeling thin biological structures [42, 43], they fall short in capturing the bending behavior present in bifurcations or high curvature regions of thin branched structures undergoing large deformations [44]. Shell formulations are particularly effective for describing the mechanics of structures with a small thickness-to-length ratio and pronounced bending. However, traditional shell formulations require additional rotational degrees of freedom to represent bending behavior and therefore come with higher computational costs [45]. We recently developed a rotation-free shell formulation for vascular structures which accounts for both membrane and bending behaviors using only displacement degrees of freedom [46]. Various shell formulations have been successfully implemented in FSI problems with IBM. For example, Gilmanov et al. [45] used a curvilinear IBM with a finite volume fluid solver and a rotation-free shell solid solver to simulate aortic valves. Nitti et al. [47] employed a direct forcing IBM with a finite difference discretization of the fluid, coupled with an isogeometric analysis-based shell solver to model interactions between thin structure and incompressible flow. Boustani et al. [48] used a sharp interface IBM with a finite difference fluid solver and a shell solid solver. However, all of these implementations rely on structured fluid grids, limiting their capacity to handle complex geometrical fluid domains, such as patient-specific vascular geometries.

Finite elements method (FEM) is a powerful tool for modeling fluid dynamics in cardiovascular systems [49, 50] using unstructured grids. Zhang et al. [51] developed an IBM model in the context of FEM, referred to as an immersed finite element method (IFEM). In this approach, finite element discretizations were employed for both fluid and solid domains, while a reproducing kernel particle method–based regularized delta function was used to transfer kinematic and force information between nonconforming meshes. Such kernel-based interpolation requires additional numerical support and quadrature operations in the coupling procedure. The structural domain was modeled using a general three-dimensional solid formulation, well-suited for thick-walled structures, but high computational cost when applied to thin structures. IFEM has been applied to simulations of red blood cells, aortic stents, and whole-heart models [52].

In this study, a partitioned IBM framework is developed to leverage efficient and specialized solvers for the fluid and solid subproblems. Two preexisting solvers are employed: CRIMSON, a stabilized FEM solver for the fluid domain [53], and a FEM–based nonlinear, rotation-free shell formulation for the thin structural domain [44, 46]. The shell formulation was specifically developed for vascular biomechanics applications and has been previously validated in aortic simulations. A direct forcing IBM [36] is adopted due to its ease of implementation within existing solid solvers and its relatively low computational cost. Fluid–structure coupling is achieved using predefined finite element shape functions for the fluid solver, which reduces the need for additional interpolation and associated computational overhead at the interface. Both weak and strong time-coupling strategies are implemented. To our knowledge, this work presents the first IBM formulation that integrates a FEM fluid solver with a rotation-free shell model to simulate interactions between thin structures and incompressible flows. The formulation is validated using two three-dimensional benchmark problems involving thin structures. A simplified valve-like model is used to illustrate the performance of weak and strong coupling strategies. Finally, the method is demonstrated in an idealized type-B aortic dissection geometry.

## 2. Methods

### 2.1. Fluid solver

The fluid phase is governed by the incompressible Navier–Stokes equations, given by

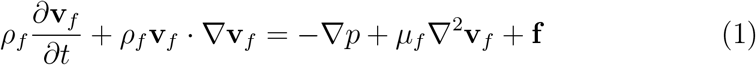

and

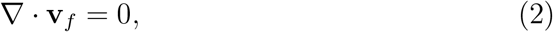

where **v**_*f*_ represents the fluid velocity, *p* is the pressure, *ρ*_*f*_ is the fluid density, *µ*_*f*_ is the dynamic viscosity, and **f** a vector of body forces. For the boundary conditions, the boundary Γ of the fluid domain Ω_*f*_ is decomposed into a Dirichlet boundary Γ_*g*_ and a Neumann boundary Γ_*h*_ such that

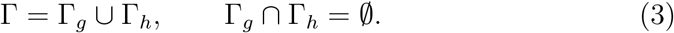

The fluid equations are discretized using a stabilized finite element formulation. Let the trial solution and weighting function spaces be defined as

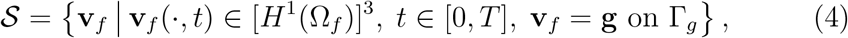

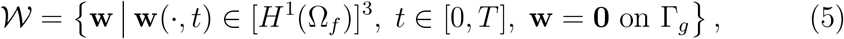

And

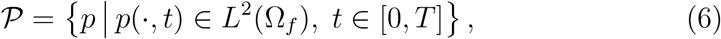

where *H*^1^(Ω_*f*_ ) denotes the Sobolev space of square-integrable functions with square integrable weak derivatives over the fluid domain Ω_*f*_, **g** is the prescribed Dirichlet boundary condition, and *L*^2^(Ω_*f*_ ) denotes the space of square-integrable functions over Ω_*f*_ .

Multiplying Eqs. (1) and (2) by these test functions, integrating over the fluid domain Ω_*f*_, applying integration by parts, and adding stabilization term yields the following weak form: find **v**_*f*_ *∈ 𝒮* and *p ∈ 𝒫* such that for every **w** *∈ 𝒲* and *q ∈ 𝒫*,

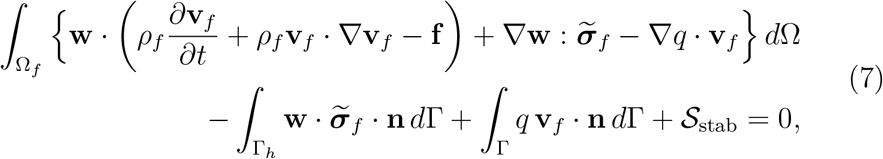

where 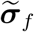 is the Cauchy stress tensor for an incompressible Newtonian fluid, defined as

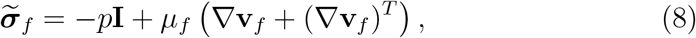

and *𝒮*_stab_ denotes the stabilization term [54, 55]:

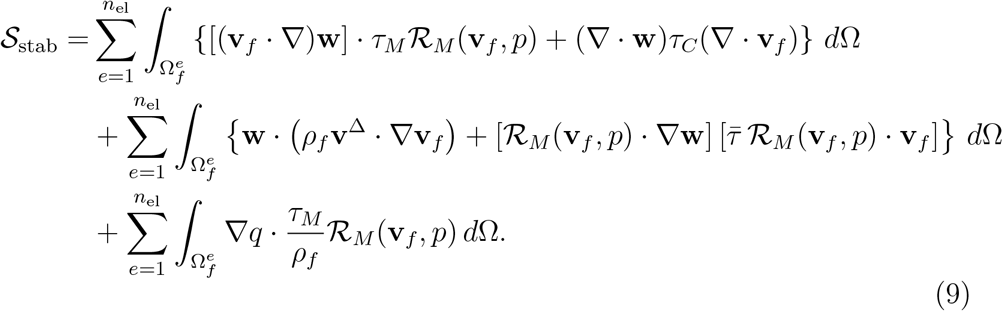

Here, *τ*_*M*_, *τ*_*C*_, and 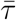 are stabilization parameters, and ℛ_*M*_ (**v**_*f*_, *p*) is the residual of the momentum equation, defined as

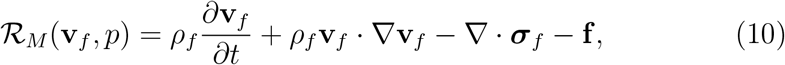

The conservation-restoring advective velocity is defined as

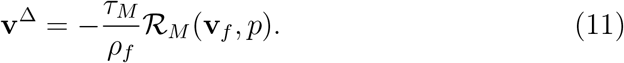

The fluid velocity and pressure fields are approximated using standard finite element linear shape functions:

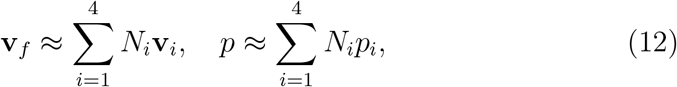

where *N*_*i*_ denotes the *i*th linear shape function associated with the three-dimensional unstructured tetrahedral elements. Equal-order linear interpolation is employed for both the velocity and pressure fields.

The resulting semi-discrete system of equations is integrated in time using the generalized-*α* method [56], which provides second-order accuracy and controlled numerical dissipation. The method treats the inertial and viscous terms implicitly, leading to unconditional stability for the fluid solver. This finite element formulation for the fluid problem is implemented within the open-source software CRIMSON [53]. The finite element mesh consists of linear tetrahedral elements. Meshing was conducted using the `MeshSim` libraries in CRIMSON [57].

### 2.2. Solid solver

Solid structures of interest in this work, such as dissection flaps and valves, have a small thickness-to-length ratio. Membrane models allow us to simplify three-dimensional structural problems by solving them on a two-dimensional middle surface with finite thickness entering parametrically in the constitutive relations, thereby greatly reducing computational cost [58, 59]. However, membrane models are not capable of capturing bending stiffness, which can lead to numerical buckling in complex geometries, such as bifurcation [60, 61, 62, 44]. Shell models can capture bending stiffness but are computationally more expensive due to the additional rotation degree of freedom. In this study, the solid problem is solved using a nonlinear rotation-free shell formulation for vascular biomechanics [46]. This formulation calculates both the membrane and bending behavior via displacement degrees of freedom for a triangular element and its three neighboring elements, making it a good fit for use in an efficient fluid-structure interaction framework. This method has been validated for large deformations [46]. The nonlinear rotation-free shell formulation uses FEM to solve an Elasto-dynamic equation via

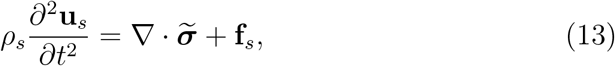

where **u**_*s*_ represents the solid displacement, *ρ*_*s*_ is the solid density, 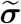 is the Cauchy stress tensor, and **f**_*s*_ is the external force, such as gravity acting as a body force and pressure or contact forces acting as surface forces.

Human aortic tissues have been considered as incompressible materials from a biomechanical perspective [63, 64]. Incompressible Neo-Hookean hyper-elastic constitutive model is implemented in this study. The strain energy density function, *W*, for this model is

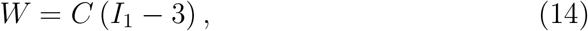

where *I*_1_ is the first principal invariant of the right Cauchy–Green deformation tensor and *C* is a material constant.

The Cauchy stress tensor 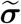 appearing in Eq. (13) is derived from the strain energy density function *W* . For an incompressible Neo-Hookean material, the constitutive relation is obtained as

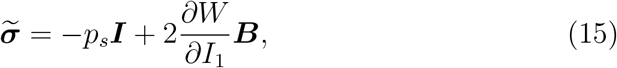

***B*** =***FF***^*T*^ is the left Cauchy–Green deformation tensor, with ***F*** denoting the deformation gradient. *p*_*s*_ is a Lagrange multiplier enforcing incompressibility.

The solid domain is discretized using two-dimensional unstructured triangular elements. Meshing is performed using `Gmsh`, an open-source finite element grid generator [65]. The generalized-*α* method is used for time integration [66]. In the generalized-*α* method, numerical dissipation is controlled by the spectral radius at infinite frequency, *ρ*_*∞*_, which appears in the definition of the algorithmic parameters *α*_*m*_ and *α*_*f*_ . Following Chung and Hulbert [67], the generalized-*α* method parameters are defined as

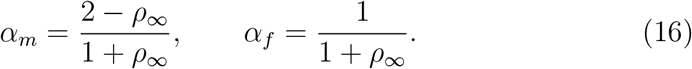

The parameters *α*_*m*_ and *α*_*f*_ determine the temporal weighting of the acceleration and internal force terms in the semi-discrete solid equations, thereby controlling numerical dissipation of the high-frequency response. Among different choices of the user-specified spectral radius at infinite frequency, *ρ*_*∞*_, the asymptotic annihilation case (*ρ*_*∞*_ = 0) is employed for the rotation-free shell formulation, resulting in maximum numerical dissipation of high-frequency response.

### 2.3. Fluid-structure interaction

The fluid and solid domains are coupled using a direct forcing immersed boundary method (IBM). In this study, we used two forces which enable the interaction between fluid and solid solvers. These forces ensure a kinematic and dynamic equilibrium between fluid and solid:

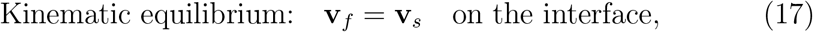

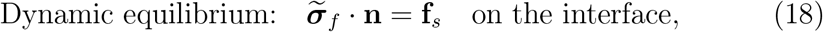

where **v**_*f*_ and **v**_*s*_ are the fluid and solid velocities, 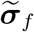 is the fluid stress tensor, **n** is the unit normal vector to the interface, and **f**_*s*_ is the external force acting on the solid. The first force is an IBM body force, which enforces the kinematic equilibrium and is applied as a body force in the Navier– Stokes equations in Eq. (1). The second force is the hydrodynamic force, which enforces dynamic equilibrium and is used as the external force in the elastodynamic equation (13). Details of these forces are described below.

#### 2.3.1. IBM body force

An IBM with direct forcing, as introduced by Uhlmann [36], is used to calculate the IBM body force. In this work, the method is further extended to support computations on unstructured grids. The direct forcing IBM enforces the no-slip condition (**v**_*f*_ = **v**_*s*_) at the interface by adding a fictitious body force at the solid nodes of the interface. The following steps describe the IBM body force calculation process:

a. The fluid elements that contain each solid node are first identified. The fluid solutions at the four fluid nodes of the tetrahedral fluid element are then interpolated onto the solid node using the FEM shape functions (Fig. 1 (a)):

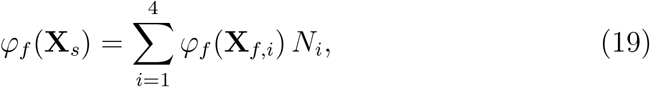

where *φ*_*f*_ denotes the fluid solution, including the velocity, pressure, and the first and second derivatives of the velocity. **X**_*s*_ and **X**_*f,i*_ are the solid and fluid node locations, respectively, and *N*_*i*_ is the *i*-th linear shape function used for the finite element method (see Eq. (12)). By employing these pre-defined FEM shape functions, additional computational costs associated with interpolation can be avoided.
b. The IBM body force is calculated at the solid node accordingn to

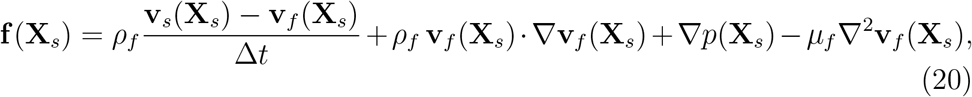

where **v**_*s*_ is the solid velocity and **v**_*f*_ is the fluid velocity interpolated from the surrounding fluid nodes.
c. The IBM body force calculated at the solid node is redistributed to the fluid nodes. This process is carried out in two steps. In the first step, the IBM body force at the solid node is interpolated to the four fluid nodes using the same FEM linear shape functions as in Eq. (19) (Fig. 1 (b)):

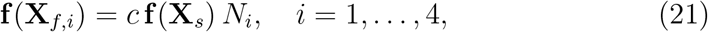

where *c* is the scaling factor used to enforce conservation of the total force acting on the fluid. The scaling factor calculation follows the method introduced by Nitti et al. [47]:

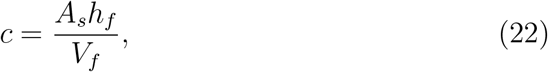

where *A*_*s*_ is the local solid mesh area, *h*_*f*_ is the local fluid mesh size, and *V*_*f*_ is the local fluid mesh volume. If more than one solid node is located near a fluid element, the IBM body force contributions from each solid node are accumulated.

**Figure 1:**
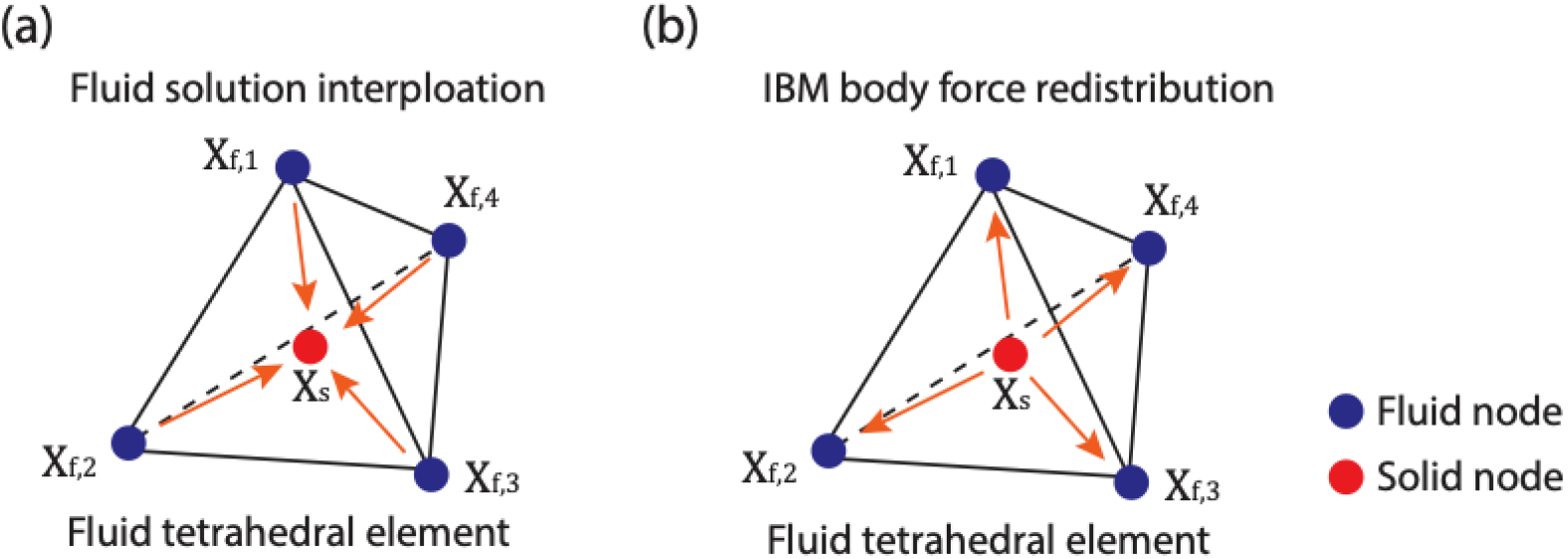
Interpolation using FEM shape functions. Interpolation (a) from fluid to solid and (b) from solid to fluid.

In the second step, we employ a diffusion equation to ensure a smooth distribution over the fluid domain. In IBM, the body force exerted by the solid is formally represented by a Dirac delta function. In numerical implementations, this singular force must be regularized and spread over a finite support of width *δ*_*f*_ to ensure numerical stability and grid-independent solutions. Traditionally, this regularization is achieved by convolving the Lagrangian forces with a compactly supported kernel function, such as a cosine kernel [31] or moving least-squares weighting functions [68], thereby distributing the force to surrounding Eulerian grid points. While effective for structured grids, these approaches become computationally expensive for unstructured meshes, as each Lagrangian marker requires identifying and communicating with a potentially large number of neighboring Eulerian cells.

To overcome this limitation, Capecelatro and Desjardins [69] proposed an Euler–Lagrange strategy in which force regularization is performed in two stages: (i) a local mollification step that transfers the force to the nearest Eulerian cell, followed by (ii) a diffusion process that spreads the force over the desired length scale *δ*_*f*_ . This approach avoids explicit support-domain searches and enables efficient force spreading on unstructured grids. Importantly, the diffusion step can be solved implicitly, yielding a robust and computationally efficient regularization procedure.

Following this strategy, after the IBM force is locally distributed to the fluid nodes using FEM shape functions, we solve the following diffusion equation for each force component to obtain a smooth body-force field on the Eulerian grid:

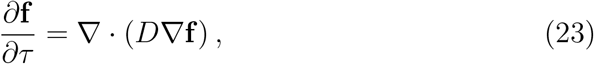

where *D* is a diffusivity and *τ* is a pseudo-time [69]. The smoothing length scale is set to 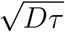 . The equation is discretized in pseudo-time using the backward Euler method and solved in a single time step with an iterative conjugate-gradient solver. The diffusion coefficient and time-step size are selected such that the diffusive length scale matches the size of the supported domain. This approach is robust and does not introduce oscillations as the immersed Lagrangian points move between Eulerian elements [70]. This method is also compatible with adaptively refined local grids. The final IBM body force is applied as the body force **f** in the fluid momentum equation (1).

#### 2.3.2. Hydrodynamic force

The hydrodynamic force acting on the open solid surface is computed following the method proposed by Nitti et al. [47]. This approach has been shown to provide accurate results and to effectively handle large pressure differences across the solid surface [71]. The method employs two probe points to calculate the hydrodynamic force [68], as illustrated in Fig. 2, where the probe points are aligned along the outward and inward normal directions of the solid surface (**n**^+^ and **n**^*−*^). For clarity, a two-dimensional triangular fluid mesh and a one-dimensional solid surface are depicted in Fig. 2. The distance from the solid node to the probe point, denoted by h, is determined based on the local fluid mesh size [72].

**Figure 2:**
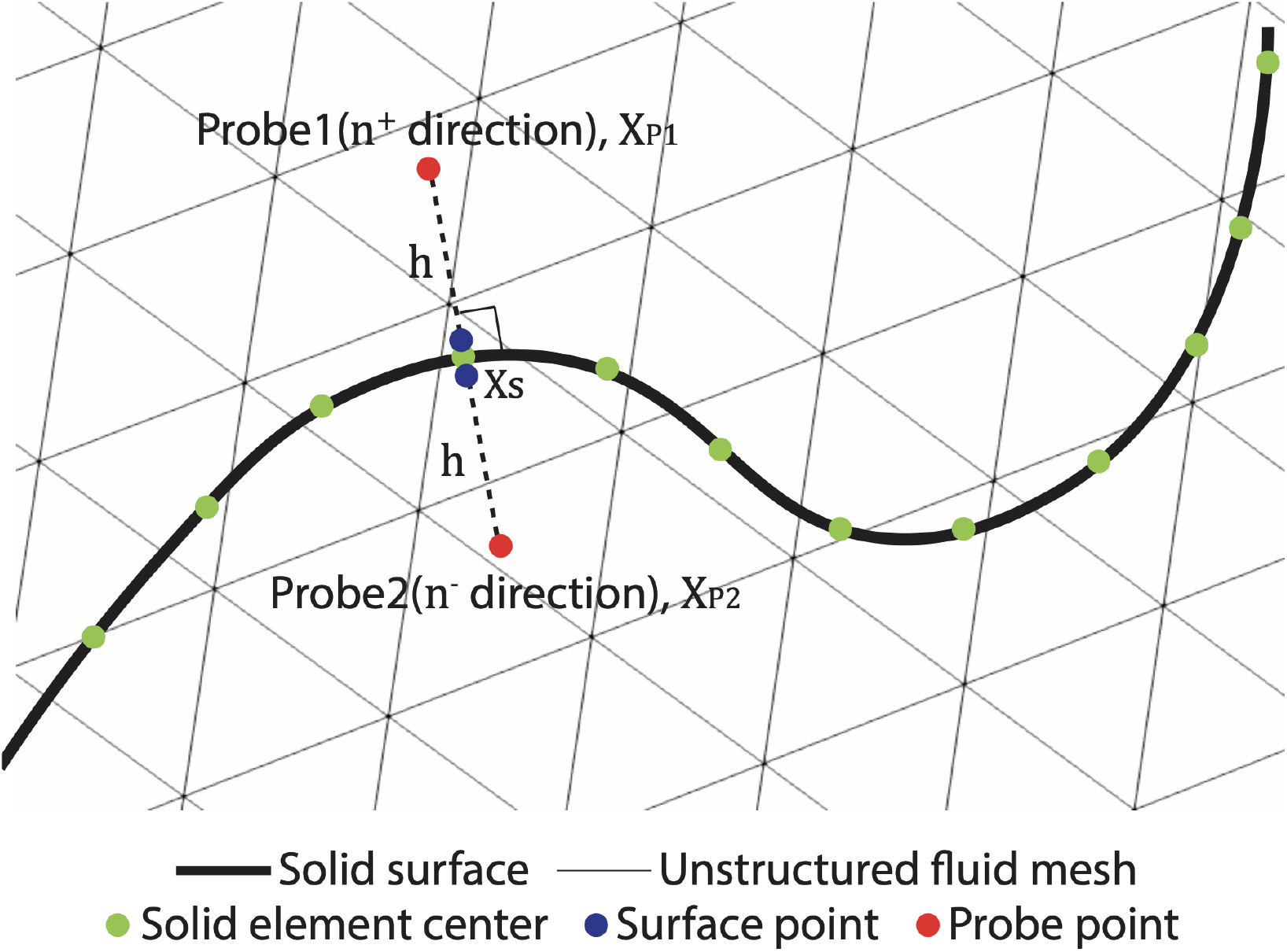
Hydrodynamic force calculation using two probe points, where h is the local fluid mesh size.

The hydrodynamic force **F**, associated with the unit normal vector **n** and the fluid stress tensor 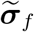, is computed as

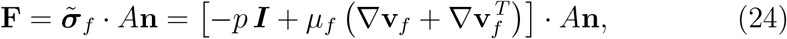

where ***I*** is the identity tensor and *A* is the area of the solid element. For an open solid surface, this force is written as

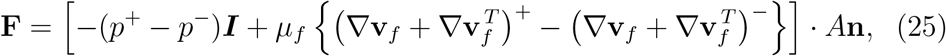

The pressure at the surface probe point *p*^+^ is computed using a boundary-layer equation derived from the momentum equation projected onto the normal direction, i.e.

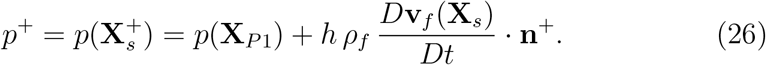

In this equation, the material derivative of the fluid velocity, *D***v**_*f*_ */Dt*, is approximated by the solid node acceleration **a**_*f*_ (**X**_*s*_) [47]. The pressure and acceleration at the probe point and the solid node are evaluated using the same interpolation procedure based on the FEM shape functions (Eq. (19)).

The fluid velocity is assumed to vary linearly in space within the boundary layer near the solid surface [73], which ensures that the first derivative of the velocity is identical at the solid node and the corresponding probe points. Under this assumption, the velocity gradient at the surface point is evaluated as

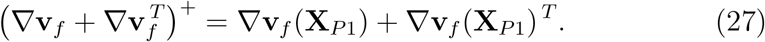

The velocity gradient at the probe point is evaluated using the same interpolation procedure based on the FEM shape functions (Eq. (19)). The resulting hydrodynamic force **F** is applied as the external force **f**_*s*_ in the solid momentum equation (13).

### 2.4. Solid searching

Searching for solid points within an unstructured grid can be computationally expensive. This section describes the procedure for tracking the Lagrangian solid points as they move within a partitioned unstructured Eulerian fluid tetrahedral mesh. At the beginning of the simulation, as a preprocessing step, the host fluid element for each solid point is identified. Although this initial search requires scanning the entire fluid mesh for each solid point and can be computationally demanding, it is performed only once.

Figure 3 illustrates a schematic of the solid tracking method. The core tracking algorithm is based on a Delaunay search, which evaluates the barycentric coordinates of a solid point relative to a fluid element [74]. If all barycentric coordinates are positive, the solid point lies within the corresponding fluid element; if any coordinate is negative, the point lies in a neighboring fluid element across the face associated with the negative coordinate. Using the neighboring connectivity of the fluid elements, the algorithm advances to the indicated adjacent element and repeats the inclusion test. The time-step size of the FSI simulation is selected to limit the motion of the immersed solid, such that solid points traverse no more than a few fluid elements per time step. As a result, the search procedure typically converges within a small number of iterations, making it reliable and efficient in practice.

**Figure 3:**
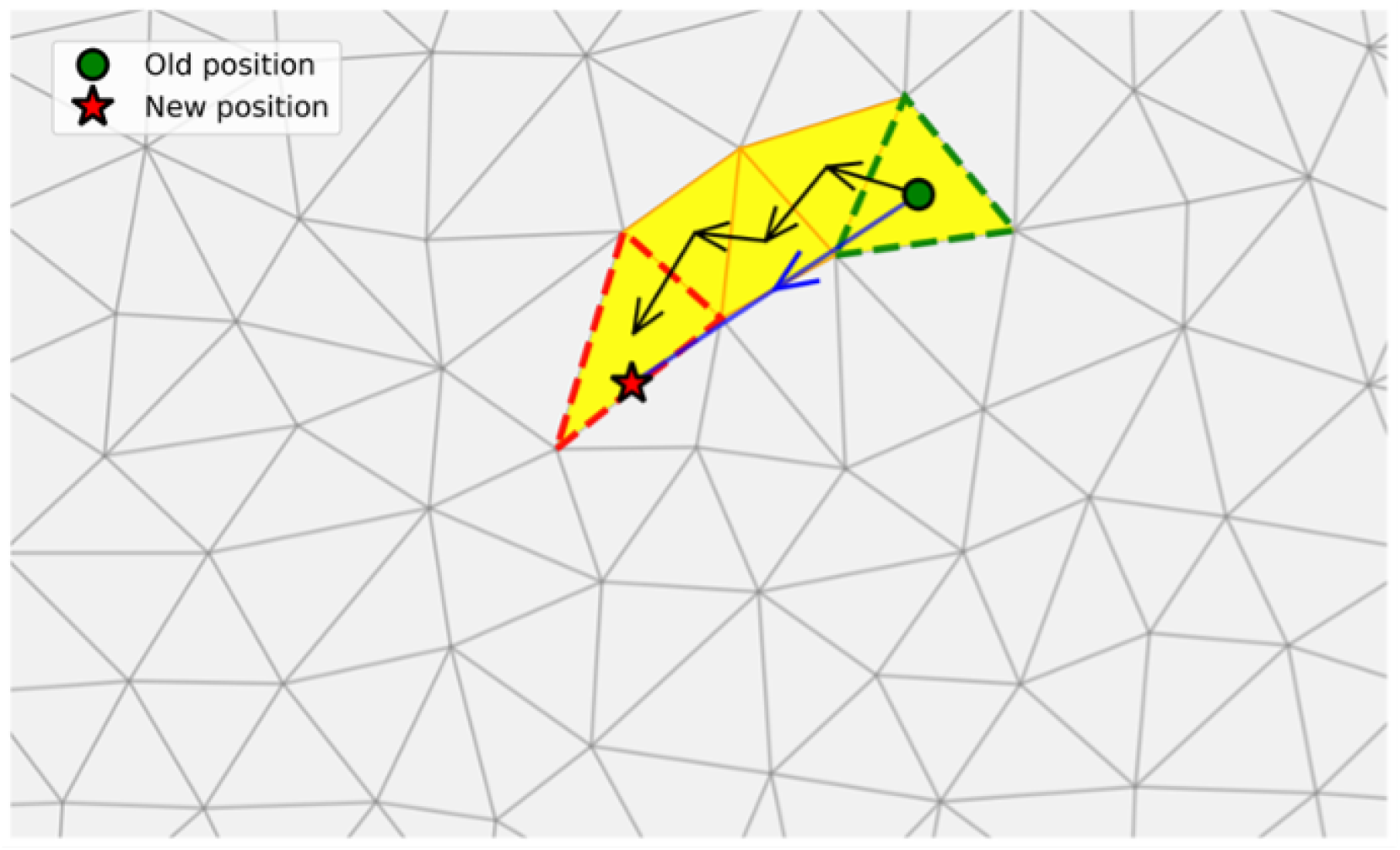
Schematic of the solid tracking method in an unstructured grid. The blue arrow denotes the motion of the solid point over one fluid time step, and the black arrows indicate the iterative tracking path. The initial and final fluid elements containing the solid point are shown in green and red, respectively.

This tracking scheme is applied independently to four sets of solid points: solid mesh nodes, two probe points, and solid element centers. The solid mesh nodes move according to the computed kinematics and are tracked as described previously. Although the solid element centers and probe points do not move independently, their positions are determined based on the updated solid mesh. As a result, they can be reliably tracked using the same procedure since their displacements remain small.

### 2.5. Time coupling scheme

A partitioned approach, in which the fluid and solid problems are solved in a staggered manner, is employed for time coupling. Two coupling strategies are considered: weak (explicit/staggered/loose) and strong (implicit/iterative) coupling [19, 75]. Weak coupling solves the fluid and solid problems once per time step and is computationally efficient, having been successfully applied in aerodynamic applications [76, 77, 78]. However, numerical instability and degradation of temporal accuracy have been reported for cases with low solid-to-fluid density ratios (*ρ*_*s*_*/ρ*_*f*_ ) [26, 79, 80]. These issues arise from the added-mass effect, in which a component of the hydrodynamic pressure force acts as an additional mass term in the solid equations [81, 82]. In contrast, strong coupling iteratively solves the fluid and solid problems within a single time step until the coupled solutions converge, thereby ensuring numerical stability and maintaining kinematic and dynamic equilibrium at the end of each time step [20, 83]. To reduce the computational cost associated with strong coupling, an under-relaxation scheme is employed [20, 26]. Both weak and strong coupling approaches are implemented in this study and are detailed below.

#### 2.5.1. Weak and strong coupling algorithms

In this section, we describe the weak and strong coupling strategies employed in this study. The overall coupling framework is formulated using a set of modular operators: the IBM body force calculation (*BF* ), the fluid solver (*FS* ), the hydrodynamic force evaluation (*HDF* ), the solid solver (*SS* ), and the under-relaxation operator (*UR*). The quantities 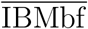 and 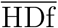 denote the IBM body force and the hydrodynamic force, respectively. The parameter *λ* is the under-relaxation coefficient, and *ϵ* = 10^*−*4^ is used as the convergence tolerance. The subscript *I* denotes intermediate quantities prior to the application of under-relaxation. Weak coupling advances the fluid and solid solvers once per time step (see Algorithm 1), whereas strong coupling employs a predictor–corrector strategy with sub-iterations (see Algorithm 2).

##### Algorithm 1

Weak coupling

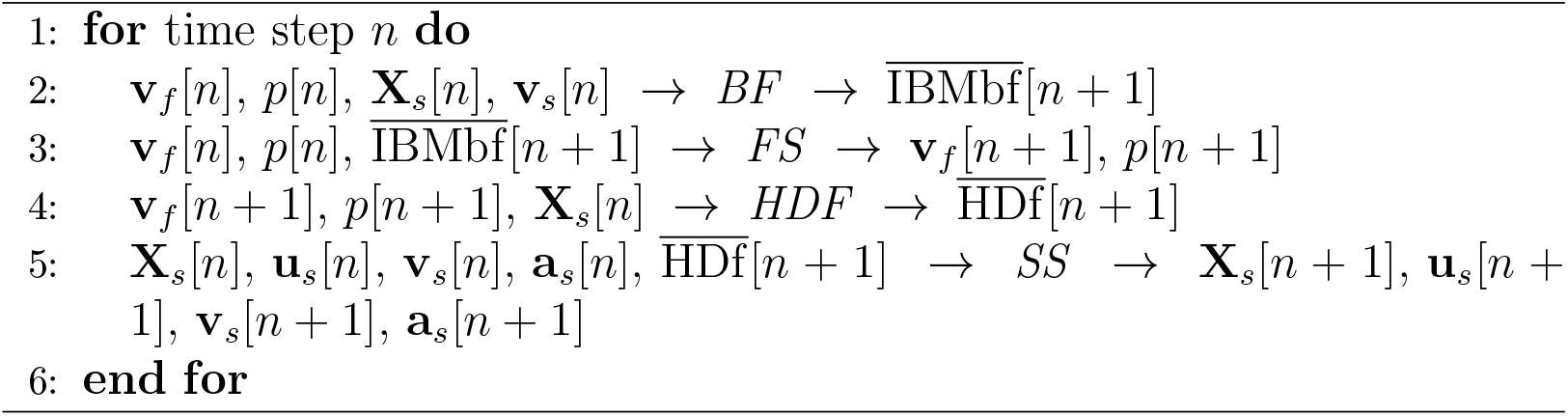

##### Algorithm 2

Strong coupling

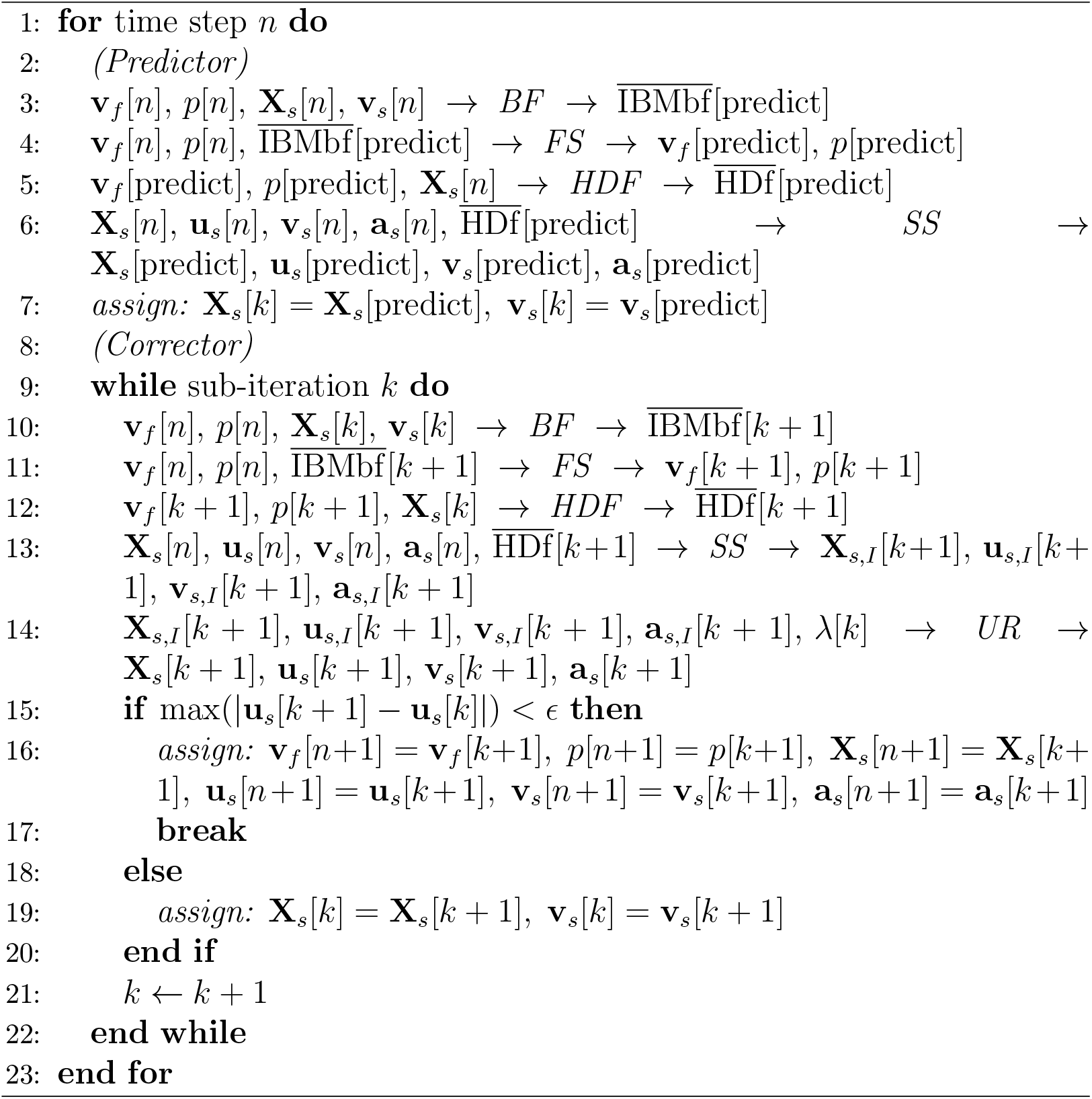

#### 2.5.2. Under-relaxation scheme

Strong coupling with iterative weak coupling can require many iterations to converge [82]. An under-relaxation scheme with Aitken acceleration [84] has been shown to reduce the number of coupling iterations per time step [20, 47, 85, 86]. In this study, we follow the under-relaxation scheme used by Nitti et al. [47]. The under-relaxation operator *UR* conducts following sub-steps:

a. *Calculate under-relaxation coefficient for the Aitken method*.

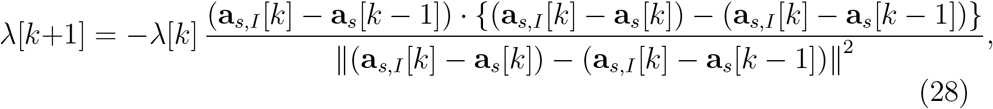

The initial value for the first time step is set to *λ*[1] = 0.3, and for subsequent time steps *λ*[1] is initialized using the final value of *λ* from the previous time step.
b. *Update acceleration with under-relaxation*.

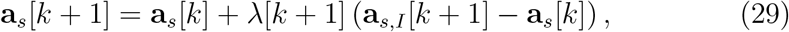
c. *Update velocity, displacement, and position using Newmark formulations*.

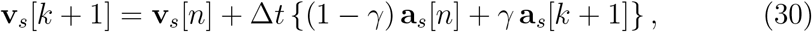

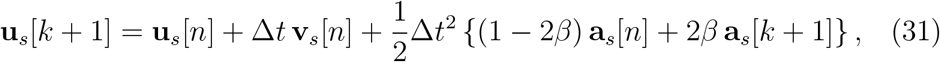

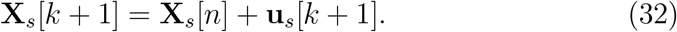

The Newmark parameters are set to *γ* = 1.5 and *β* = 1.0. Typically, with this under-relaxation scheme, two to four coupling iterations are sufficient to achieve convergence within a single time step.

## 3. Results and discussion

In this paper, we consider four benchmark problems. The first problem is designed to validate the computation of hydrodynamic forces, while the second focuses on assessing the FSI capabilities of the proposed framework. The third problem investigates strong and weak time-coupling schemes using a simplified aortic valve model. Finally, an idealized type-B aortic dissection problem is examined.

### 3.1 Validation case 1: Drag on the circular disk

Accurate evaluation of hydrodynamic forces acting on open solid surfaces is a key component of the proposed algorithm. To validate the hydrodynamic force computation, we consider the benchmark problem of a circular disk immersed in a uniform flow, as illustrated in Fig. 4. A two-dimensional rigid disk of radius *R* is fixed at the origin inside a cylindrical fluid domain extending from *−*5*R* to 30*R* in the streamwise direction (*x* direction) and having a radius of 12*R*. The fluid domain is discretized using 11.7 million uniform, unstructured tetrahedral elements with an element size of 0.02*R*, while the solid disk is discretized using 500 uniform triangular elements with the same element size, 0.02*R*. These configuration and mesh size are consistent with two previous numerical studies by Shenoy and Kleinstreuer [87] and Tian et al. [88]. Uniform inflow velocity is prescribed at the inlet, a no-slip boundary condition is imposed on the lateral wall, and zero pressure is enforced at the outlet (Fig. 4). The drag coefficient of the disk is evaluated over a range of Reynolds numbers and compared with experimental and computational results from the literature. The drag coefficient of the disk, *C*_*D*_, and Reynolds number, *Re*, are defined as

**Figure 4:**
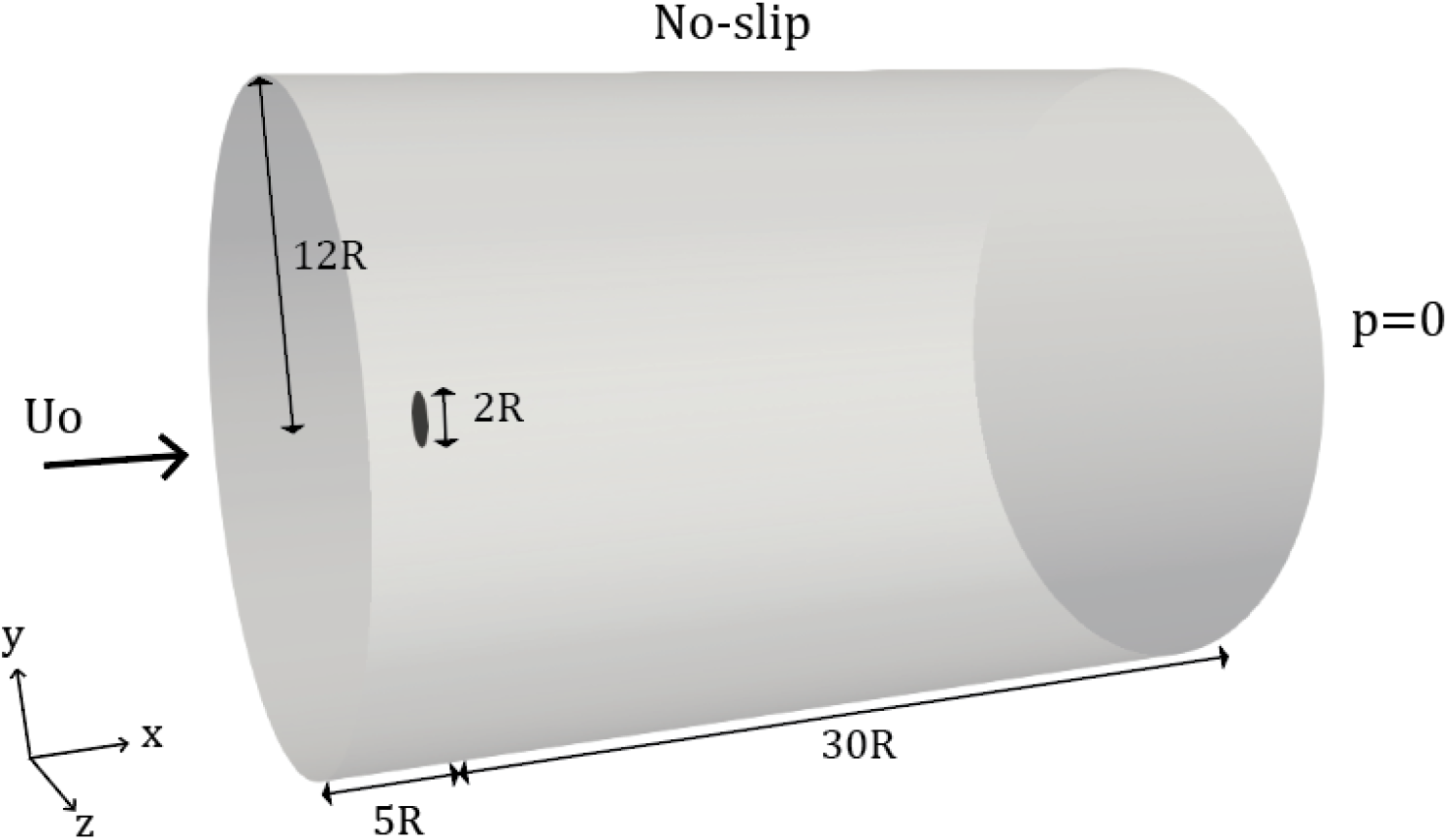
Schematic of the computational domain of the circular disk immersed in a cylinder.

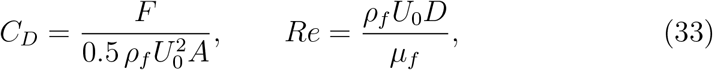

where *ρ*_*f*_ is the fluid density, *U*_0_ is the uniform inflow velocity, *A* = *πR*^2^ is the disk area, *D* = 2*R* is the disk diameter, and *µ*_*f*_ is the fluid viscosity. The hydrodynamic force acting on the disk is denoted by *F* . Seven Reynolds numbers are considered: *Re* = 10, 21, 36, 54, 75, 90, and 107. Different Reynolds numbers are obtained by varying the fluid density while keeping the inflow velocity and viscosity fixed. The drag coefficient is evaluated under fully developed flow conditions.

Figure 5 presents the drag coefficient of the disk over a range of Reynolds numbers from *Re* = 10 to 110. Reynolds numbers exceeding 110 are not considered because the flow becomes unsteady and does not reach a fully developed state [89]. The results obtained in the present study are compared with experimental measurements reported by Roos and Willmarth [89], as well as numerical simulations by Shenoy and Kleinstreuer [87], who employed a finite-volume-based solver using CFX-10.0 (ANSYS Inc., 2005) and by Tian et al. [88] using Cartesian-grid-based IBM using a cell-centered projection scheme, which is implemented on their in-house solver. All reference studies consider a finite disk thickness of 0.2*R*. Additionally, Tian et al. provide results for the two-dimensional disk with zero thickness. In this study, the nonlinear rotation-free shell formulation uses a two-dimensional mid-surface solid domain with zero thickness. The predicted drag coefficients show good agreement with the numerical results corresponding to zero disk thickness across the entire Reynolds number range.

**Figure 5:**
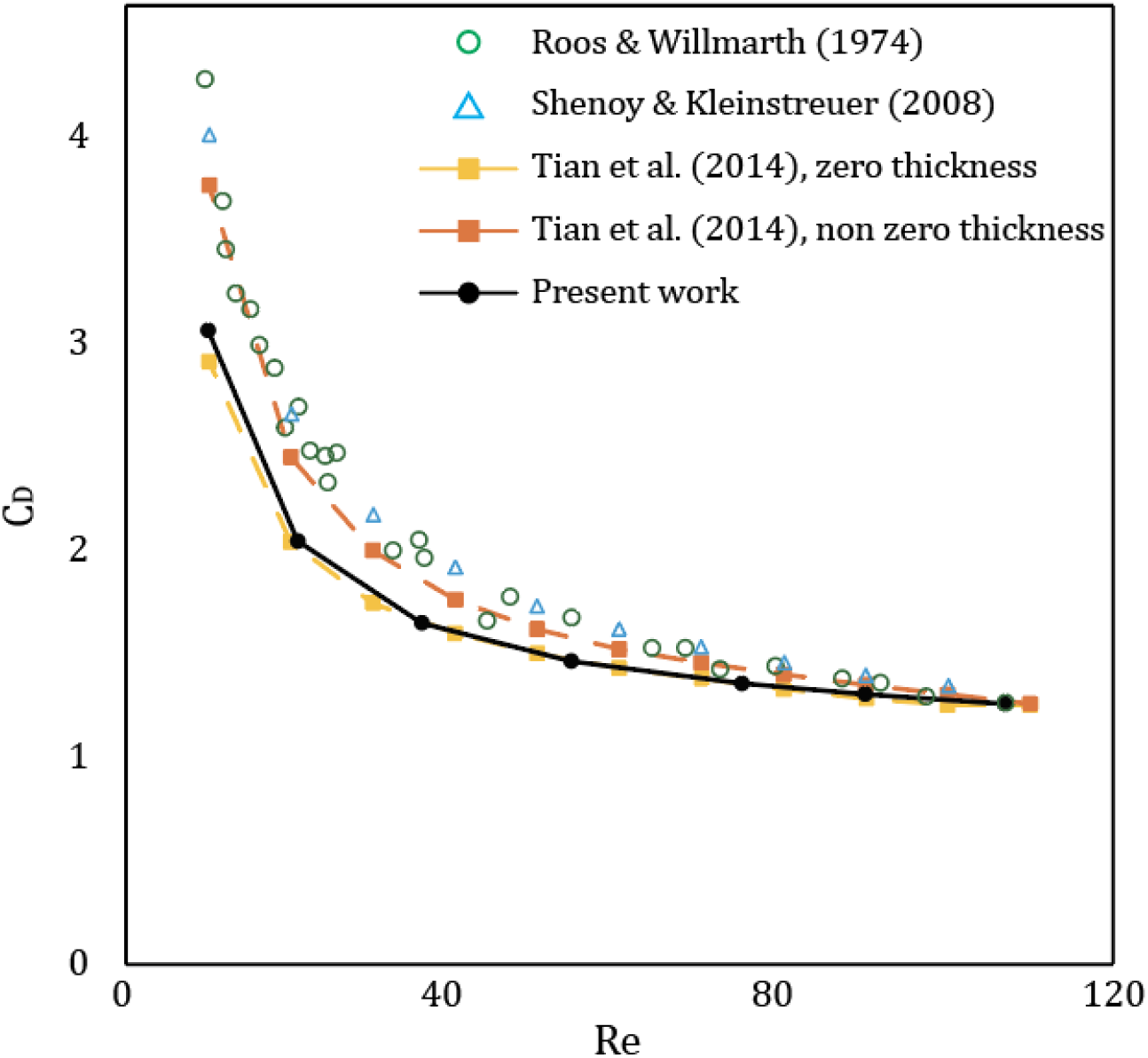
Comparison of drag coefficient on a circular disk for various Reynolds numbers.

### 3.2. Validation case 2: Three-dimensional flapping flag in a uniform flow

This section considers a flapping flag in a uniform flow problem, originally proposed by Huang and Sung [90]. This problem has been widely used as a benchmark for evaluating IBM algorithms due to the large three-dimensional structural displacements involved [47, 88]. Figure 6(a) illustrates the computational domains of this problem. A square flag of side length *L* is placed inside a cylindrical fluid domain, with the center of the leading edge located at the origin. The cylindrical domain extends from *−*2*L* to 8*L* in the stream-wise direction (*y*-direction) and has a radius of 5*L*. The fluid domain is discretized using 2.1 M nonuniform, unstructured tetrahedral elements, with a minimum element size of 0.02*L* near the flag (see Fig. 6(b)); the nonuniform mesh is generated using the mesh adaptation feature in CRIMSON. The solid domain consists of 5.8k uniform triangular elements with an element size of 0.02*L*. This mesh configuration is consistent with two previous numerical studies by Huang and Sung [90] and Tian et al. [88]. The fluid boundary conditions include a uniform inflow at the inlet, a uniform velocity condition on the lateral wall, and zero pressure at the outlet (Fig. 6(a)). The initial condition corresponds to a uniform flow with velocity *U*_0_ in the streamwise direction (*y*-direction). The leading edge of the flag is fixed, while the remaining three edges are free to move. Initially, the flag is tilted by an angle of 0.1*π* in the *y −z* plane, as shown in Fig. 6(b).

**Figure 6:**
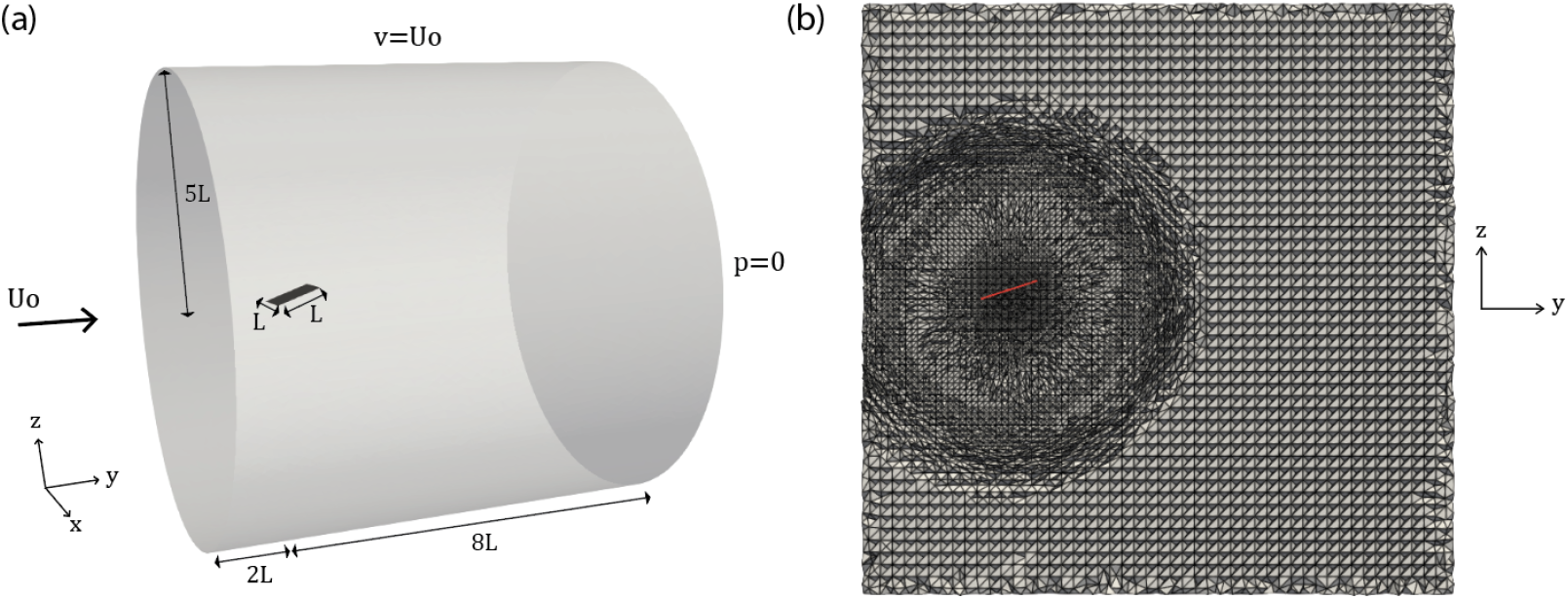
(a) Schematic of the computational domain of flapping flag in a cylinder and (b) nonuniform, unstructured fluid mesh (*x* = 0 slice).

The Reynolds number is set to *Re* = 200. The mass ratio between the fluid and the solid is defined as 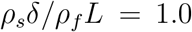, where *δ* denotes the flag thickness used in the rotation-free shell formulation. The nondimensional bending rigidity is defined as

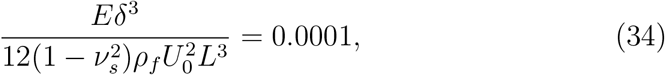

where *E* and *v*_*s*_ are the elastic modulus and Poisson’s ratio of the solid, respectively. An incompressible Neo-Hookean constitutive model is employed, Poisson’s ratio is set to *v*_*s*_ = 0.5. The time-step size is chosen as Δ*t* = 0.002*L/U*_0_ to ensure that the CFL number remains below 1.0.

Figure 7(a) shows the velocity contours at *y − z* plan (*x* = 0 slice), illustrating the disturbed flow field induced by the motion of the flag. Figure 6(b) presents the time history of the transverse (*y*-directional) displacement of the midpoint at the trailing edge of the flag. The results from the present study are compared with data from two previous computational studies by Huang and Sung [90] and Tian et al. [88]. Good agreement is observed between the present results and the reference data, despite the use of different constitutive models in the previous studies, e.g., Saint Venant–Kirchhoff model.

**Figure 7:**
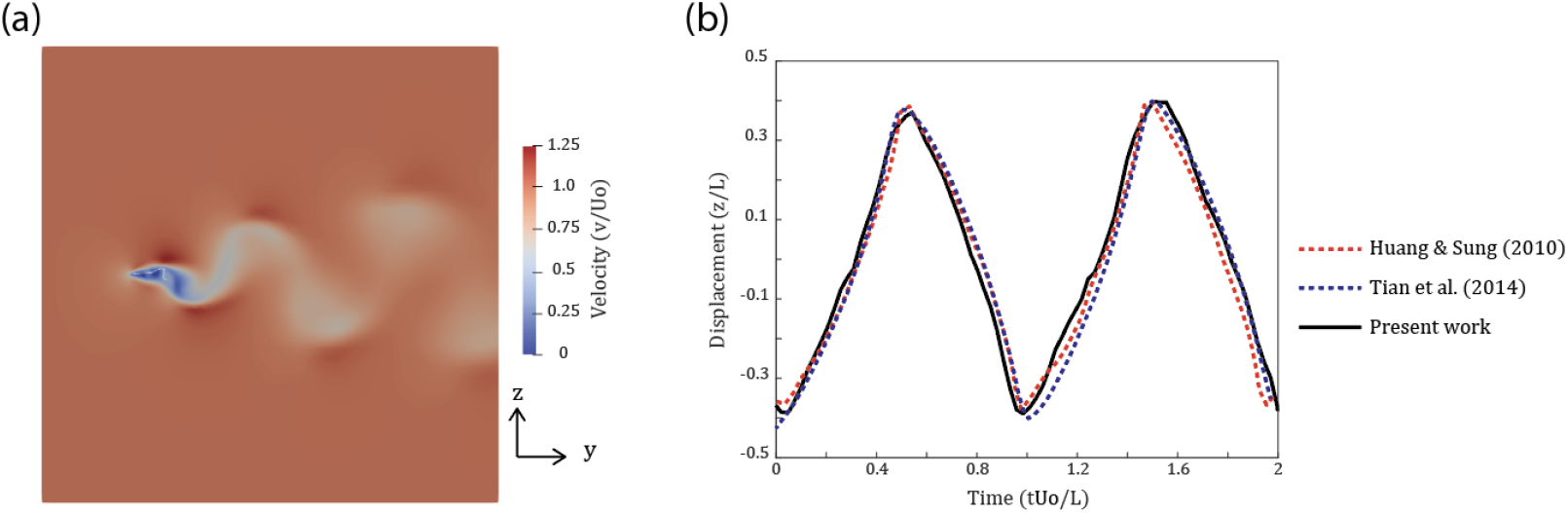
(a) Velocity contour at the *y − z* plane (*x* = 0 slice) and (b) the transverse (*z*-directional) displacement of the midpoint at the trailing edge of the flag

### 3.3. Strong and weak coupling comparison

The problem of a thin vertical obstacle immersed in a cylindrical tube (see Fig. 8) is considered to compare strong and weak time coupling schemes. The solid-to-fluid density ratio, *ρ*_*s*_*/ρ*_*f*_, has been identified as a key parameter in selecting an appropriate coupling strategy, with strong coupling generally recommended for cases where *ρ*_*s*_*/ρ*_*f*_ *<* 1 [26, 79, 80]. To investigate this effect, four density ratios, *ρ*_*s*_*/ρ*_*f*_ = 0.5, 1, 10, and 100, are examined by varying the solid density while keeping the fluid density constant at 0.00106 g*/*mm^3^. The fluid viscosity is set to 0.004 g*/*mm·s.

**Figure 8:**
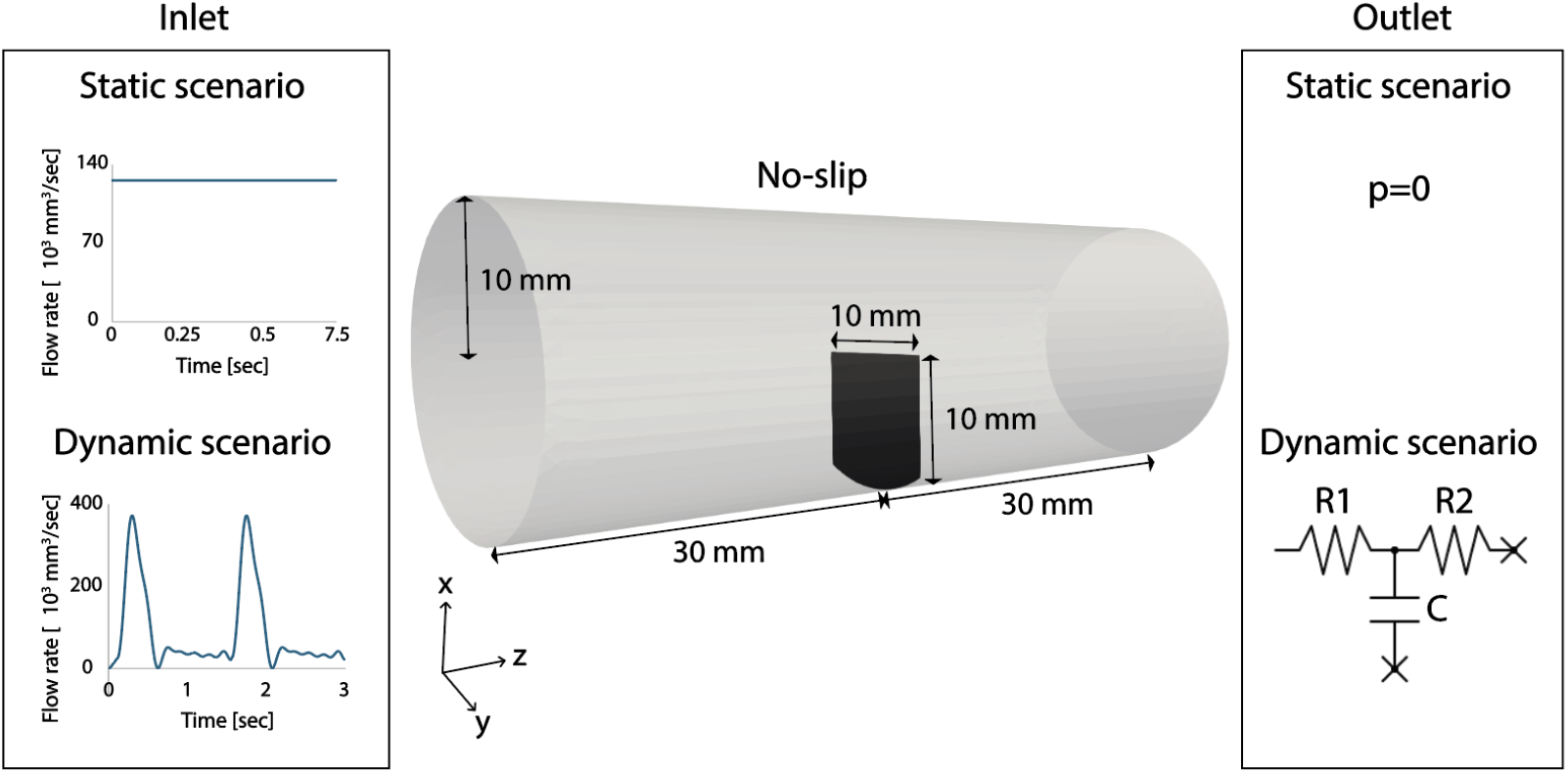
Schematic of the computational domain for the problem of a thin vertical obstacle immersed in a cylindrical tube and boundary conditions for the static (upper boundary conditions) and dynamic (lower boundary conditions) scenarios.

Both static and dynamic scenarios are considered to assess the performance of the weak and strong coupling schemes under different physical conditions. In static problems, the solution reaches an equilibrium state and remains unchanged over time, whereas in dynamic problems, the solution continuously evolves. The computational domain is identical for both scenarios and presents in Fig. 8. The vertical solid structure, with a width of 10 mm, is located at the middle of the cylindrical domain. The structure has a thickness of 0.5 mm. The bottom edge was fixed while the remaining three edges are free to move. The cylindrical domain has a radius of 10 mm and a length of 60 mm.

Mesh independence studies are conducted using four meshes with characteristic element sizes of 1 mm, 0.7 mm, 0.5 mm, and 0.3 mm. These correspond to 120k, 360k, 970k, and 3.2M fluid elements, and 150, 370, 1, 000, and 2, 500 solid elements, respectively. Convergence of the solid displacement at the final time step is achieved, with a maximum difference of 0.7% between the 0.5 mm and 0.3 mm meshes. Based on this result, all subsequent simulations are performed using the 0.5 mm mesh size. With this configuration, the fluid domain consists of 970k uniform, unstructured tetrahedral elements, while the solid domain consists of 1, 000 uniform triangular elements.

#### 3.3.1. Static scenario

First, the static scenario is considered. A constant inflow condition and a zero-pressure boundary condition are prescribed at the inlet and outlet of the fluid domain, respectively (see Figure 8). The time-step size is set to ∆*t* = 0.005 s to ensure that the CFL number remains below 1.0. The simulation is performed for 1, 500 time steps, corresponding to a total simulation time of 7.5 s. The average Reynolds number is 2, 000, and the elastic modulus of the solid is set to 10^4^ Pa.

Figure 9 presents the velocity contour on the *x − z* plane (*y* = 0 slice) at the final time step for *ρ*_*s*_*/ρ*_*f*_ = 1 using the strong coupling scheme. As the fluid moves toward the streamwise direction (*z*-direction), the solid structure correspondingly deforms in that same direction.

**Figure 9:**
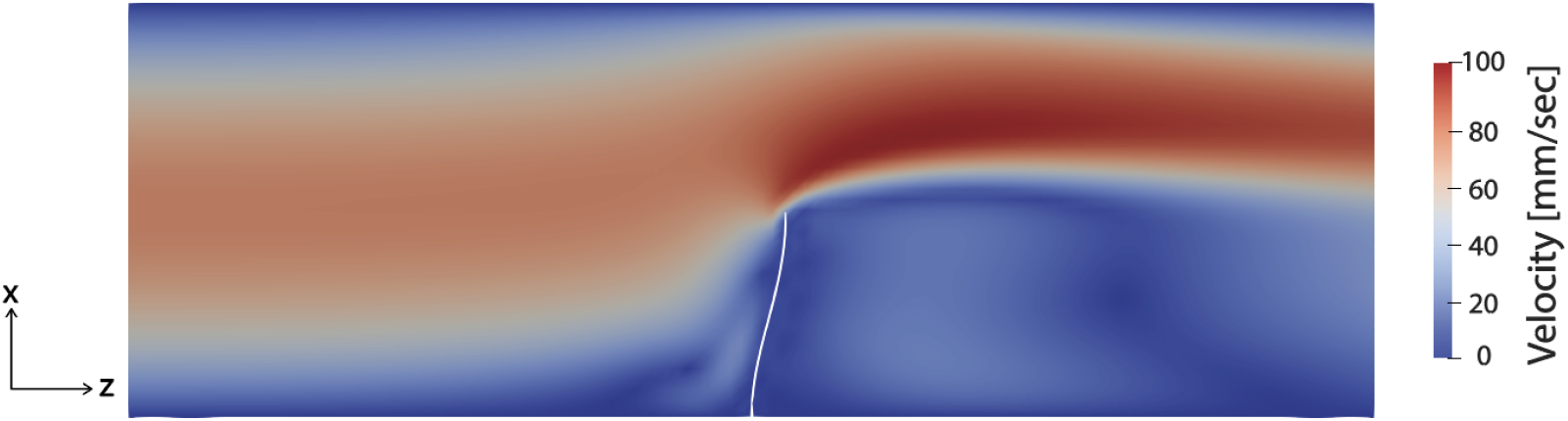
Velocity contour on the *x − z* plane (y=0 slice) of static scenario with *ρ*_*s*_*/ρ*_*f*_ = 1 using strong coupling

Figure 10 presents the time history of the *z*-directional displacement of the midpoint on the top edge of the solid structure. Panels (a),(b),(c), and (d) compare the displacement results obtained using the weak and strong coupling schemes for solid-to-fluid density ratios of *ρ*_*s*_*/ρ*_*f*_ = 0.5, 1, 10, and 100, respectively. In these plots, the red and black curves correspond to the strong and weak coupling results, respectively.

**Figure 10:**
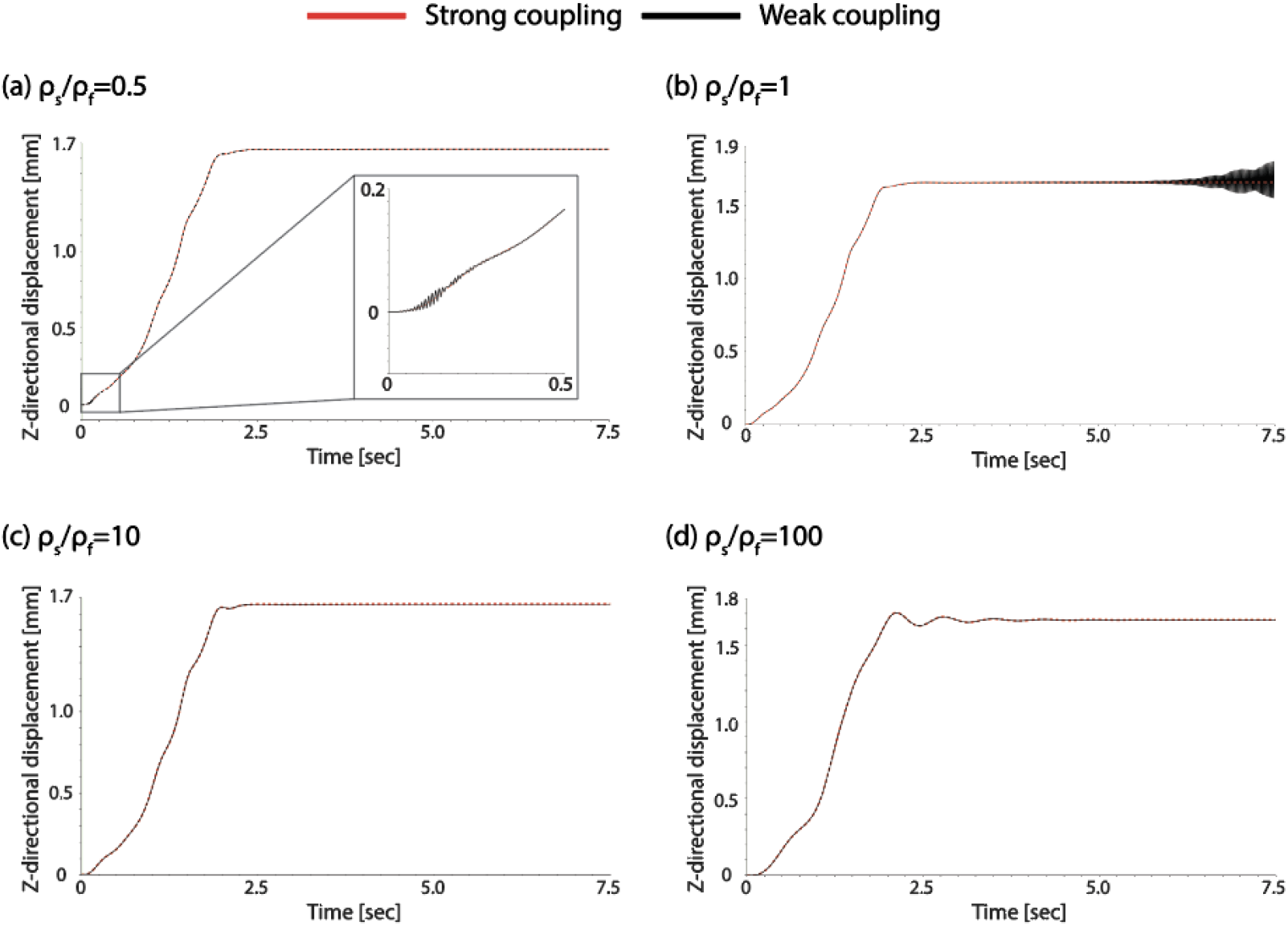
Static scenario. Strong vs weak coupling comparison with four different solid-to-fluid density ratios: (a) 0.5, (b) 1, (c) 10, and (d) 100. Red line: strong coupling and black line: weak coupling results.

For lower solid-to-fluid density ratios, the weak coupling scheme exhibits nonphysical oscillations, appearing at early times for *ρ*_*s*_*/ρ*_*f*_ = 0.5 and at later times for *ρ*_*s*_*/ρ*_*f*_ = 1. In contrast, the strong and weak coupling results show good agreement for larger density ratios (*ρ*_*s*_*/ρ*_*f*_ = 10 and 100). These observations are consistent with previous studies [26, 79, 80] and emphasize the necessity of strong coupling for problems involving low solid-to-fluid density ratios (*ρ*_*s*_*/ρ*_*f*_ *≤* 1).

Table 1 compares the computational cost of the static scenario simulations. All simulations were performed using 36 processor cores. The table reports both the total computational time for the FSI simulations and the time spent exclusively on the solid solver. In addition, the ratio of total computational time between the strong and weak coupling schemes is provided. Overall, the strong coupling scheme requires only 1.1 to 1.7 times more computational time than the weak coupling scheme. This is because, for the static problem, the solution reaches a steady state after approximately 2.5 s, beyond which no further coupling iterations are required in the strong coupling approach.

**Table 1:** Computational time for static scenarios using both strong and weak time coupling schemes with four different solid-to-fluid density ratios. Total: overall FSI simulation time. Solid: time spent solely on the solid simulation. Total and solid-only computational times are reported in units of 10^3^ s.

| Density ratio ($\rho_s/\rho_f$) | 0.5 | 1 | 10 | 100 |
| --- | --- | --- | --- | --- |
| Weak coupling (total) | 2.87 | 4.07 | 2.94 | 3.18 |
| Weak coupling (solid) | 0.21 | 0.38 | 0.21 | 0.30 |
| Strong coupling (total) | 4.84 | 4.58 | 3.96 | 4.60 |
| Strong coupling (solid) | 0.39 | 0.40 | 0.30 | 0.44 |
| Strong / weak (total) | 1.69 | 1.12 | 1.35 | 1.45 |

For the weak coupling scheme, the case with *ρ*_*s*_*/ρ*_*f*_ = 1 requires more computational time than the other cases due to oscillations occurring at later time steps. The case with the smallest density ratio (*ρ*_*s*_*/ρ*_*f*_ = 0.5) exhibits the largest ratio of total computational time between the strong and weak coupling schemes, indicating that additional coupling iterations are required to achieve convergence.

#### 3.3.2. Dynamic scenario

Next, we considered the dynamic scenario. Pulsatile flow and three-element Windkessel boundary conditions were assigned at the inlet and outlet of the fluid domain, respectively (see Figure 8). The Windkessel parameters were set to *R*_1_ = 0.02 g*/*mm^4^ s, *R*_2_ = 0.17 g*/*mm^4^ s, and *C* = 66 mm^4^ s^2^*/*g. Two cycles of pulsatile flow were simulated. The time-step size was set to ∆*t* = 5 *×*10^*−* ;4^ s to ensure that the maximum CFL number remained below 1. The simulation consisted of 6,000 time steps, corresponding to a final simulation time of 3 s. The maximum and average Reynolds numbers were 5,000 and 1,000, respectively. The elastic modulus of the solid was set to 3*×*10^5^ Pa.

Figure 11 presents the same comparison as in Figure 10, but for the dynamic scenario. The case with a large density ratio (*ρ*_*s*_*/ρ*_*f*_ = 100) was more sensitive to inlet flow fluctuations (see the inlet boundary condition in Figure 8 for the dynamic scenario), exhibiting oscillatory motion, particularly between 0.5 and 1.5 s. Unlike the static scenario, strong and weak coupling results show good agreement across all solid-to-fluid density ratios. This finding is consistent with the study by Borazjani et al. [20], who showed that strong and weak coupling yield identical results in heart valve simulations involving complex cardiac flow.

**Figure 11:**
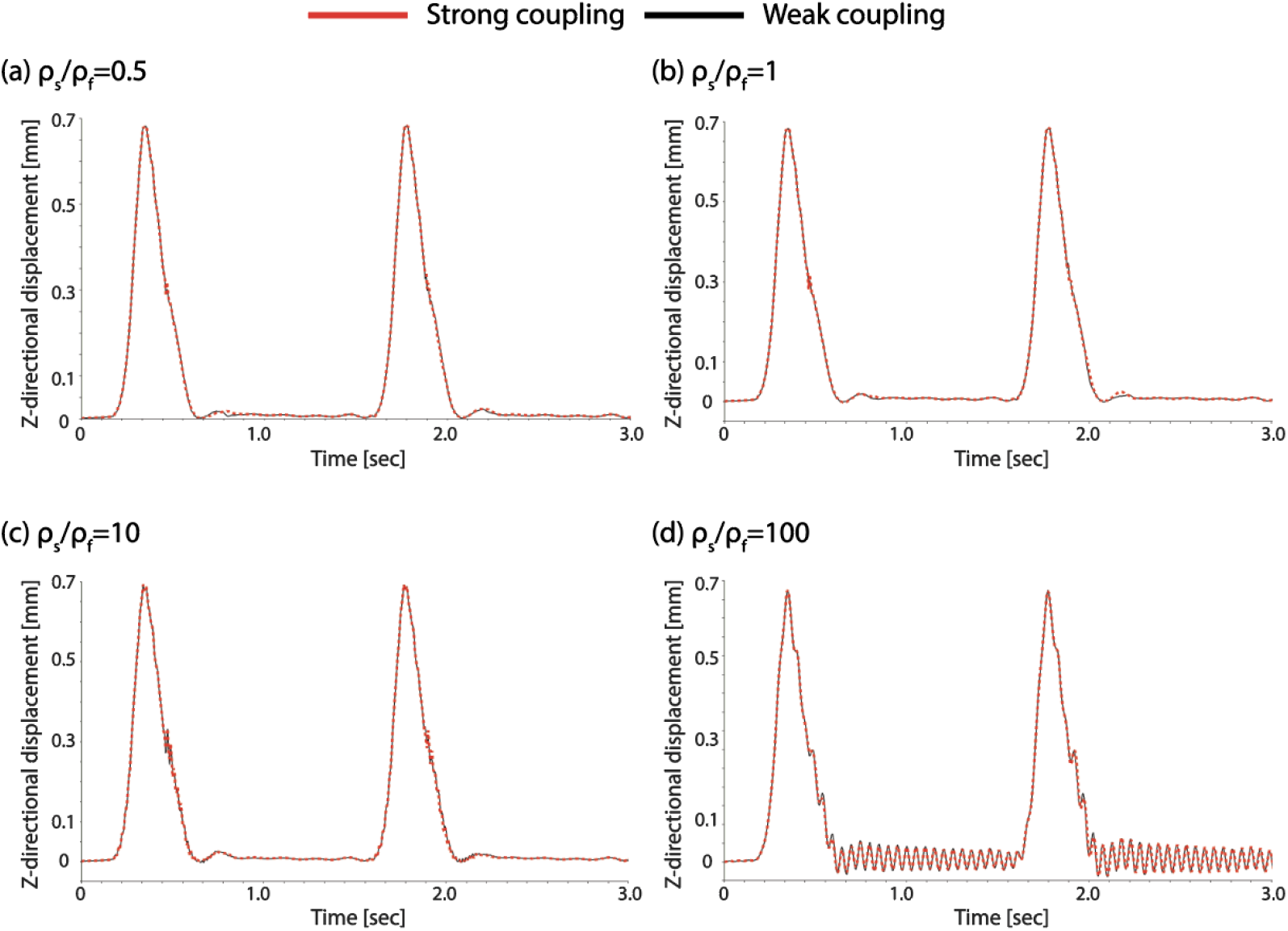
Dynamic scenario. Strong vs weak coupling comparison with four different solid-to-fluid density ratios: (a) 0.5, (b) 1, (c) 10, and (d) 100. Red line: strong coupling and black line: weak coupling results.

Table 2 presents the same data as Table 1, but for the dynamic simulation. All simulations were performed using 36 cores. Overall, the strong coupling scheme required approximately 2.1 to 2.8 times more computational time than the weak coupling scheme. Similar to the static simulations, the case with a small density ratio (*ρ*_*s*_*/ρ*_*f*_ = 0.5) exhibited a larger ratio of total computational time between the strong and weak coupling schemes, indicating that more coupling iterations were required.

**Table 2:** Computational time for dynamic scenarios using both strong and weak time coupling schemes with four different solid-to-fluid density ratios. Total: overall FSI simulation time. Solid: time spent solely on the solid simulation. Total and solid-only computational times are reported in units of 10^3^ s.

| Density ratio ($\rho_s/\rho_f$) | 0.5 | 1 | 10 | 100 |
| --- | --- | --- | --- | --- |
| Weak coupling (total) | 2.53 | 3.06 | 2.54 | 2.56 |
| Weak coupling (solid) | 0.098 | 0.099 | 0.101 | 0.102 |
| Strong coupling (total) | 7.10 | 6.45 | 5.70 | 5.19 |
| Strong coupling (solid) | 0.264 | 0.238 | 0.219 | 0.209 |
| Strong / weak (total) | 2.80 | 2.10 | 2.24 | 2.02 |

#### 3.3.3. Preference of coupling schemes

In partitioned FSI solvers, the choice of coupling scheme requires a careful balance between computational cost and numerical stability. Our results indicate that weak coupling is suitable for problems with high solid-to-fluid density ratios (*ρ*_*s*_*/ρ*_*f*_ *>* 1) in both static and dynamic scenarios. In contrast, for low density ratios (*ρ*_*s*_*/ρ*_*f*_*≤* 1), weak coupling leads to non-physical oscillations, consistent with previous observations [19, 81, 82]. In such cases, strong coupling effectively suppresses numerical instabilities, with a higher computational cost.

For dynamic problems, our results further suggest that weak coupling can remain stable over a wide range of density ratios; however, when the problem involves complex structural motion, weak coupling may still trigger instability and eventual divergence [20]. Based on these observations, strong coupling was employed for the subsequent idealized aortic dissection example. Overall, we recommend using weak coupling for dynamic problems when numerical stability is maintained, while strong coupling should be adopted when instability or divergence is observed.

### 3.4. Idealized three-dimensional type-B aortic dissection FSI model

In this example, an idealized three-dimensional geometrical model of a type-B aortic dissection is constructed (Fig. 12(a)). The fluid domain represents the aorta with a uniform diameter of 30 mm and a length of 225 mm. The solid domain corresponds to the intimal flap, as shown in Figure 12(b). The intimal flap has a uniform thickness of 1 mm and a total length of 200 mm, with one circular entry tear of 20 mm diameter and one circular exit tear of 10 mm diameter. The flap exhibits a constant curvature along the axial direction (*x*-direction) and initially divides the descending thoracic and abdominal aorta into a true lumen (TL) and a false lumen (FL), with cross-sectional areas of 218 mm^2^ and 489 mm^2^, respectively (see Figure 12(c)). The distances between the intimal flap and the aortic wall are 9 mm for the TL and 21 mm for the FL.

**Figure 12:**
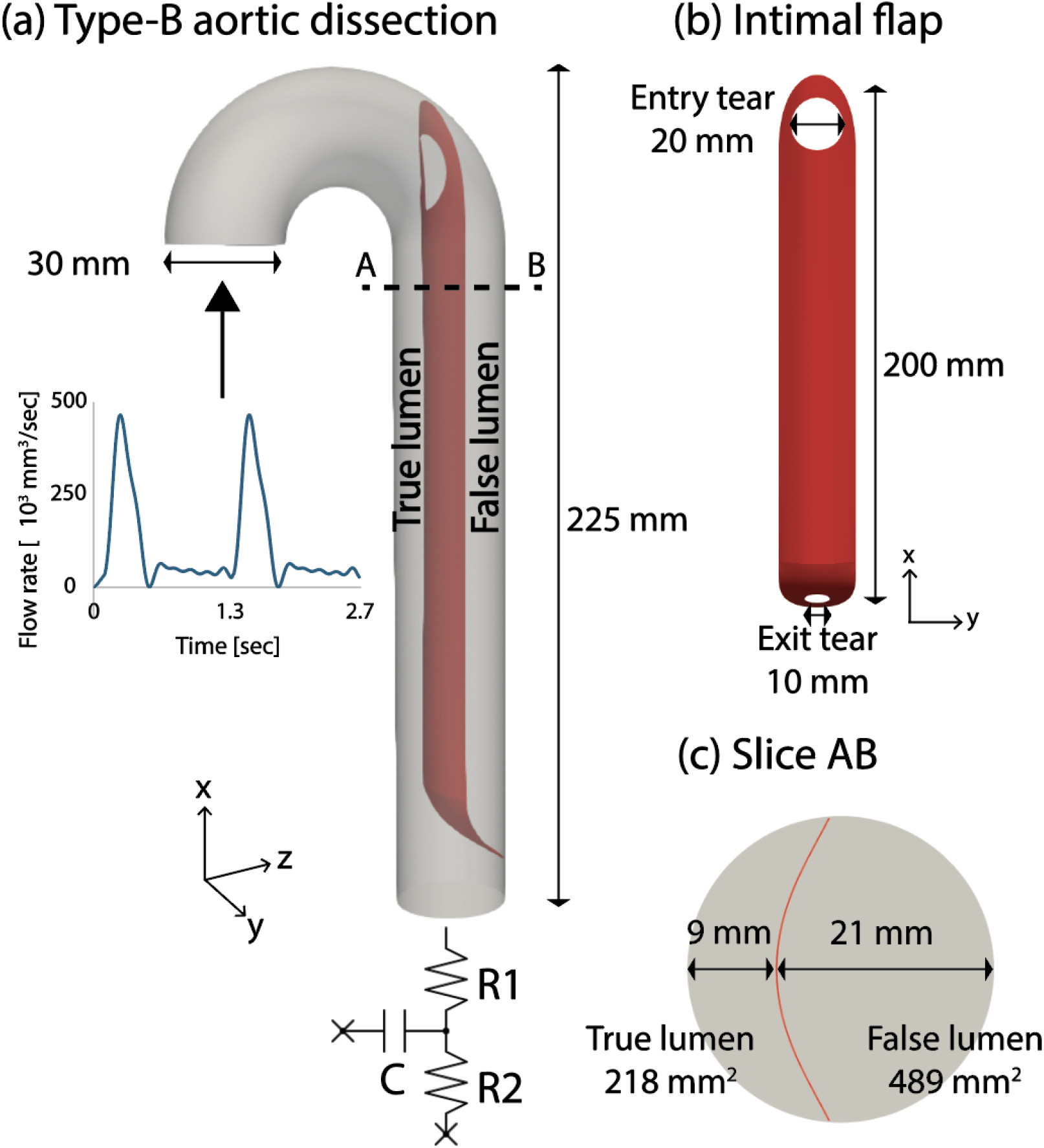
Schematic of the computational domain and boundary conditions for the idealized aortic dissection model. (a) Fluid and solid three-dimensional domain and boundary conditions. (b) Detailed solid geometry. (c) y-z slice (A-B slice).

Mesh-independence studies are conducted using three mesh sizes of 1 mm, 0.7 mm, and 0.5 mm, corresponding to 1.2M, 3.6M, and 9.5M fluid elements, respectively. Convergence is assessed based on the maximum intimal flap displacement during systole, with a maximum difference of 0.98% between the 0.7 mm and 0.5 mm meshes. All subsequent simulations are therefore performed using the 0.7 mm mesh. The fluid domain consisted of 3.6M uniform, unstructured tetrahedral elements, while the solid domain comprised 29K uniform triangular elements.

Prescribed pulsatile inflow, no-slip wall, and three-element Windkessel boundary conditions are applied at the inlet, lateral wall, and outlet of the fluid domain, respectively (Figure 12(a)). Two cardiac cycles are used for the simulation. The Windkessel parameters were set to *R*_1_ = 0.023 g*/*mm^4^ s, *R*_2_ = 0.091 g*/*mm^4^ s, and *C* = 16.54 mm^4^ s^2^*/*g. The blood density and dynamic viscosity are set to 0.00106 g*/*mm^3^ and 0.004 g*/*mm s, respectively. For the solid domain, the external edges attached to the aortic wall are fixed, while the tear edges were allowed to move freely. The density of the intimal flap is assumed to be equal to that of blood, 0.00106 g*/*mm^3^. The material stiffness of the intimal flap has not been well established in the literature. In this study, the intimal flap is assumed to be less stiff than the aortic wall, based on its anatomical origin from the inner layer of the aortic wall. This inner layer consists primarily of elastin fibers, which are more compliant than the collagen fibers that dominate the outer layers of the aortic wall [91]. Reported values for the elastic modulus of the aorta range from 0.5 to 1 *×* 10^6^ Pa [92]. Accordingly, the elastic modulus of the intimal flap is set to 2*×* 10^4^ Pa.

The time step size is chosen as ∆*t* = 1 *×*10^*−* ;4^ s to ensure that the maximum CFL number remained below 1. The simulation consisted of 27,000 time steps, corresponding to a total simulated time of 2.7 s. The maximum and average Reynolds numbers are 5,400 and 1,200, respectively.

Figure 13 illustrates the displacement of the intimal flap at diastole and systole. Panel (a) shows the three-dimensional configurations of the intimal flap. As the inlet flow increases during systole, the intimal flap deforms toward the FL, corresponding to the positive *z*-direction. The maximum displacement of the intimal flap reaches 9.79 mm and occurs near the entry tear.

**Figure 13:**
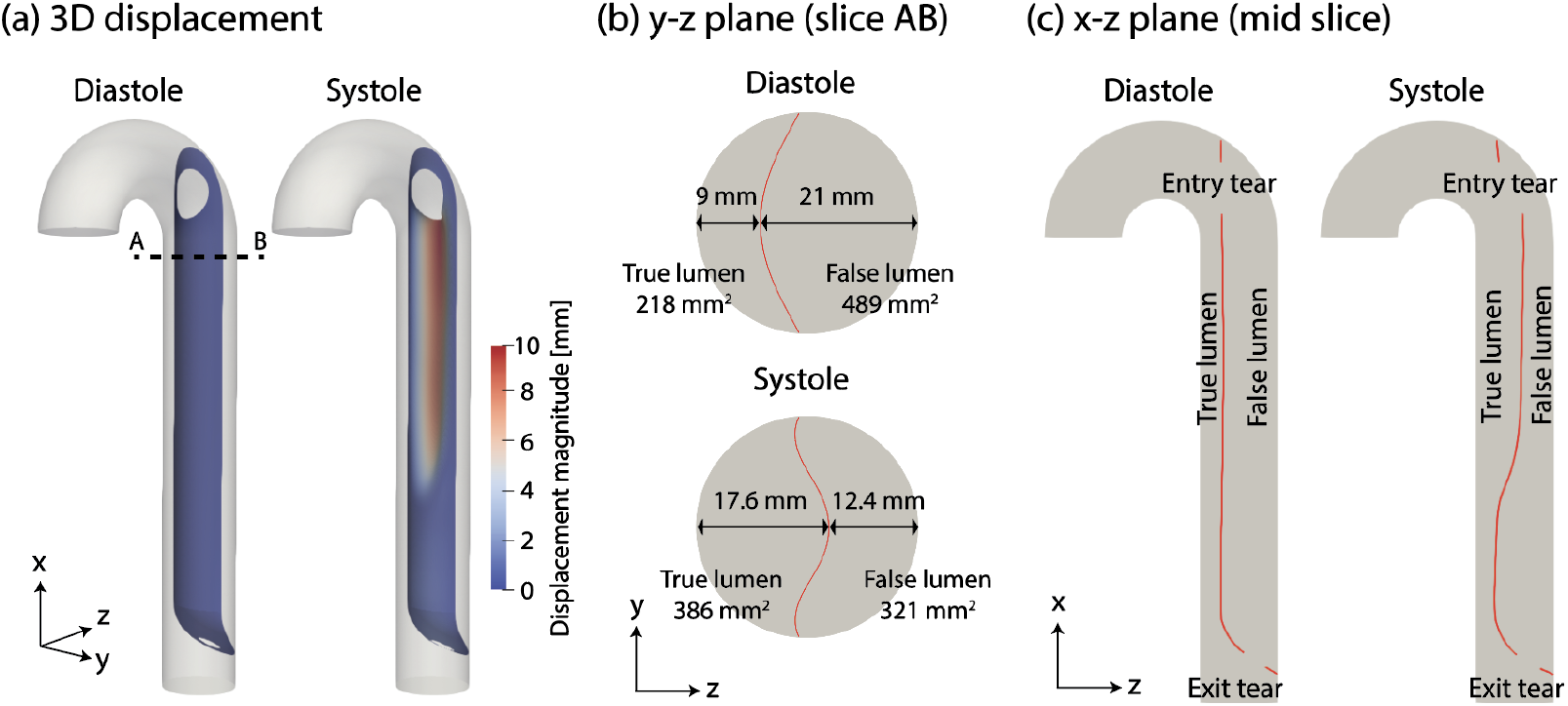
Intimal flap displacement on diastole and systole. (a) Three-dimensional displacement, (b) *y − z* plane (slice AB), and (c) *x − z* plane (mid slice)

Panel (b) presents the *y− z* plane at slice AB, located 10 mm downstream of the entry tear. At diastole, the maximum distances between the intimal flap and the aortic wall are 9 mm and 21 mm for the TL and FL, respectively.

The corresponding cross-sectional areas of the TL and FL are 218 mm^2^ and 489 mm^2^. During systole, as the intimal flap deforms, the maximum distances between the intimal flap and the aortic wall change to 17.6 mm and 12.4 mm for the TL and FL, respectively. The corresponding cross-sectional areas become 386 mm^2^ for the TL and 321 mm^2^ for the FL. Panel (c) shows the configuration of the intimal flap in the *x− z* plane at the mid-slice location. Figure 14 shows the velocity and pressure contours in different planes during diastole and systole. Panels (a) and (b) illustrate the velocity distributions on several *y− z* plane slices and on the mid *x − z* plane slice, respectively. During diastole, a relatively high velocity of approximately 140 mm/s is observed at the exit tear. This elevated velocity is attributed to the smaller cross-sectional area of the exit tear (78.5 mm^2^) compared to that of the entry tear (314 mm^2^). In addition, near the distal region of the intimal flap, the velocity in the true lumen (TL, *∼*70 mm/s) is higher than that in the false lumen (FL, *∼*50 mm/s), which is consistent with the smaller TL cross-sectional area in this region.

**Figure 14:**
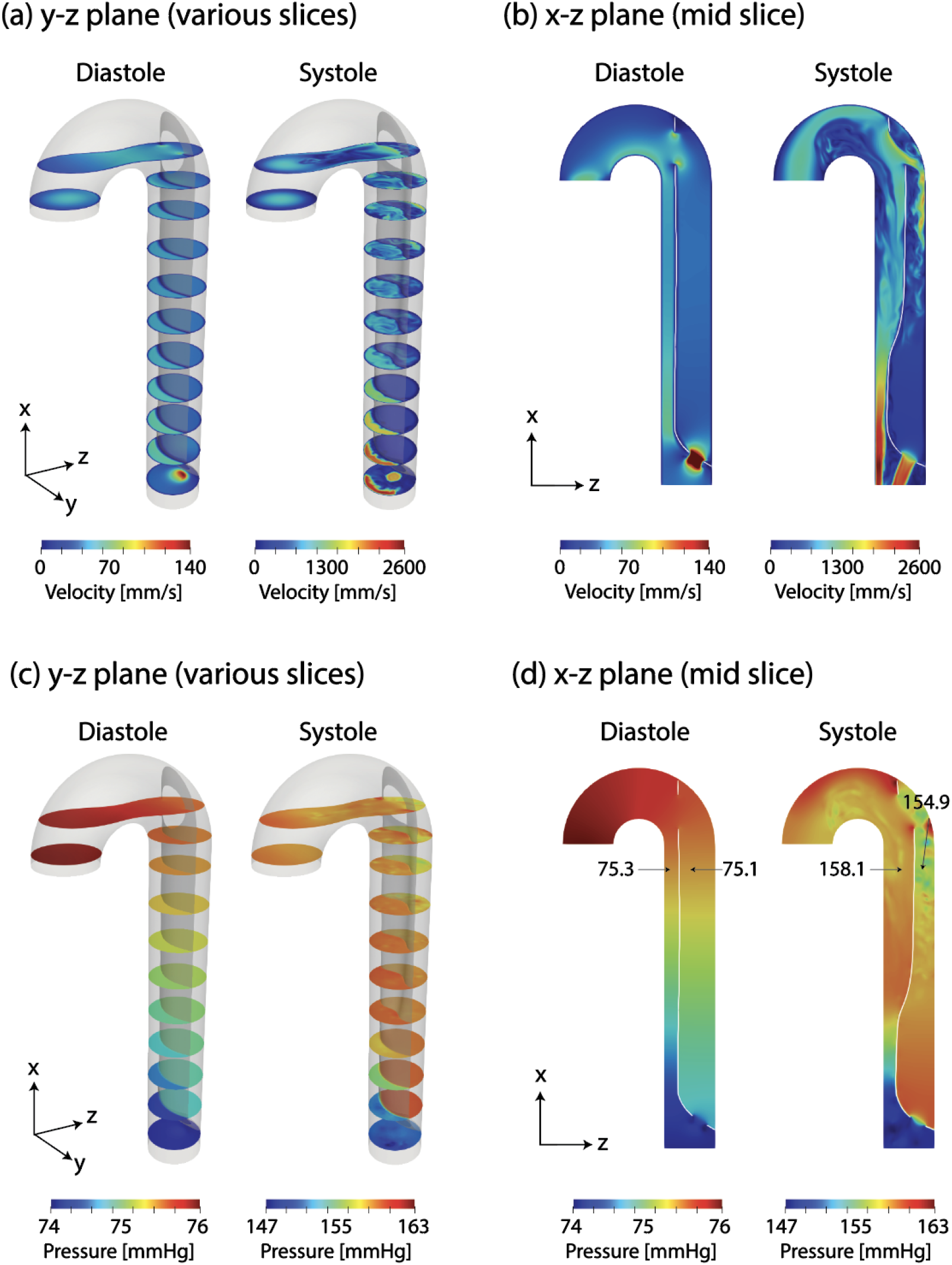
Velocity and pressure contour on the diastole and systole. Velocity: (a) *y − z* plane (various slices) and (b) *x − z* plane (mid slice). Pressure: (c) *y − z* plane (various slices) and (d) *x − z* plane (mid slice).

During systole, the increased inflow significantly raises the velocity through the entry tear to approximately 1500 mm/s, generating a jet directed toward the FL-side aortic wall (see Figure 14(b), systole). The velocity through the exit tear also increases markedly, reaching approximately 2000 mm/s. As the intimal flap moves toward the FL, the proximal TL cross-sectional area increases. Consequently, the TL velocity in this region exhibits a moderate increase to approximately 1000 mm/s. In contrast, the distal TL velocity increases sharply to approximately 2600 mm/s, where the intimal flap motion remains minimal. During systole, the large and rapid motion of the intimal flap (within less than 0.5 s) induces pronounced flow disturbances near the proximal region of the intimal flap in both the TL and FL.

Figure 14 panels (c) and (d) illustrate the pressure distributions on various *y − z* plane slices and on the mid *x − z* plane slice, respectively. The blood pressure increases from approximately 75 mmHg during diastole to 150 mmHg during systole. During diastole, the pressure difference between the true lumen (TL) and the false lumen (FL) remains below 1 mmHg. During systole, this pressure difference increases to approximately 4 mmHg, leading to large intimal flap motion in the proximal region.

These pressure results are consistent with our previous findings using an aortic dissection FSI model, which demonstrated that even relatively small pressure differences between the TL and FL are sufficient to induce significant intimal flap motion [30].

One limitation of this simplified model is that it includes only a single exit tear. In clinical cases of aortic dissection, multiple exit tears are often present. Blood flow through these additional tears can substantially alter the hemodynamics of aortic dissection, particularly the pressure difference between the TL and FL [93].

## 4. Conclusion

In this study, we developed an IBM formulation for cardiovascular simulations that couples two solvers within a direct-forcing IBM framework: FEM fluid solver using an unstructured grid and a finite element–based nonlinear rotation-free shell formulation for thin structures. Both strong and weak time-coupling schemes were implemented and applied to static and dynamic fluid–structure interaction problems over a wide range of fluid-to-solid density ratios.

By employing predefined shape functions in the fluid finite element discretization, the proposed approach minimizes the need for additional interpolation operations at the fluid–solid interface, thereby reducing computational overhead. To the best of our knowledge, this work represents the first IBM formulation that integrates a finite element fluid solver with a rotation-free shell solver, providing a robust framework for simulating interactions between thin structures and incompressible flow in complex geometries.

The proposed method was validated using two three-dimensional benchmark problems involving thin structures. As a demonstration of its applicability to clinically relevant problems, the method was further applied to an idealized type-B aortic dissection model, highlighting its potential for future cardiovascular FSI simulations. The proposed method was implemented within the open-source software CRIMSON [53], a FEM-based software designed for cardiovascular simulations.

## Notes

### Competing Interest Statement

The authors have declared no competing interest.

## References

[1] G. J. Perry, F. Helmcke, N. C. Nanda, C. Byard, B. Soto, Evaluation of aortic insufficiency by doppler color flow mapping, Journal of the American College of Cardiology 9 (4) (1987) 952–959.

[2] R. T. Eberhardt, J. D. Raffetto, Chronic venous insufficiency, Circulation 130 (4) (2014) 333–346.

[3] G. M. Deeb, H. J. Patel, D. M. Williams, Treatment for malperfusion syndrome in acute type A and B aortic dissection: a long-term analysis, The Journal of Thoracic and Cardiovascular Surgery 140 (6) (2010) S98– S100.

[4] C. J. Bennett, J. J. Maleszewski, P. A. Araoz, CT and MR imaging of the aortic valve: radiologic-pathologic correlation, Radiographics 32 (5) (2012) 1399–1420.

[5] R. Gottlob, R. May, Venous valves: morphology, function, radiology, surgery, Springer Science & Business Media, 2012.

[6] R. R. Baliga, C. A. Nienaber, E. Bossone, J. K. Oh, E. M. Isselbacher,U. Sechtem, R. Fattori, S. V. Raman, K. A. Eagle, The role of imaging in aortic dissection and related syndromes, JACC: Cardiovascular Imaging 7 (4) (2014) 406–424.

[7] H. Jilaihawi, M. Chen, J. Webb, D. Himbert, C. E. Ruiz, J. Rodés-Cabau, G. Pache, A. Colombo, G. Nickenig, M. Lee, et al., A bicuspid aortic valve imaging classification for the tavr era, JACC: Cardiovascular Imaging 9 (10) (2016) 1145–1158.

[8] H. Baumgartner, J. Hung, J. Bermejo, J. B. Chambers, T. Edvardsen,S. Goldstein, P. Lancellotti, M. LeFevre, F. Miller Jr, C. M. Otto, et al., Recommendations on the echocardiographic assessment of aortic valve stenosis: a focused update from the European Association of Cardiovascular Imaging and the American Society of Echocardiography, European Heart Journal-Cardiovascular Imaging 18 (3) (2017) 254–275.

[9] C. Nienaber, R. Spielmann, Y. Von Kodolitsch, V. Siglow, A. Piepho,T. Jaup, V. Nicolas, P. Weber, H. Triebel, W. Bleifeld, Diagnosis of thoracic aortic dissection. Magnetic resonance imaging versus transesophageal echocardiography., Circulation 85 (2) (1992) 434–447.

[10] D. Spinelli, F. Benedetto, R. Donato, G. Piffaretti, M. M. Marrocco-Trischitta, H. J. Patel, K. A. Eagle, S. Trimarchi, Current evidence in predictors of aortic growth and events in acute type B aortic dissection, Journal of Vascular Surgery 68 (6) (2018) 1925–1935.

[11] C. J. François, M. Markl, M. L. Schiebler, E. Niespodzany, B. R. Landgraf, C. Schlensak, A. Frydrychowicz, Four-dimensional, flow-sensitive magnetic resonance imaging of blood flow patterns in thoracic aortic dissections, The Journal of Thoracic and Cardiovascular Surgery 145 (5) (2013) 1359–1366.

[12] J. J. Westenberg, S. D. Roes, N. Ajmone Marsan, N. M. Binnendijk,J. Doornbos, J. J. Bax, J. H. Reiber, A. De Roos, R. J. van der Geest, Mitral valve and tricuspid valve blood flow: accurate quantification with 3D velocity-encoded MR imaging with retrospective valve tracking, Radiology 249 (3) (2008) 792–800.

[13] Y. Bazilevs, K. Takizawa, T. E. Tezduyar, Computational fluid-structure interaction: methods and applications, John Wiley & Sons, 2013.

[14] G. Hou, J. Wang, A. Layton, Numerical methods for fluid-structure interaction—a review, Communications in Computational Physics 12 (2) (2012) 337–377.

[15] B. Hübner, E. Walhorn, D. Dinkler, A monolithic approach to fluid– structure interaction using space–time finite elements, Computer Methods in Applied Mechanics and Engineering 193 (23-26) (2004) 2087– 2104.

[16] C. Michler, S. Hulshoff, E. Van Brummelen, R. De Borst, A monolithic approach to fluid–structure interaction, Computers & Fluids 33 (5-6) (2004) 839–848.

[17] D. Kamensky, M.-C. Hsu, D. Schillinger, J. A. Evans, A. Aggarwal,Y. Bazilevs, M. S. Sacks, T. J. Hughes, An immersogeometric variational framework for fluid–structure interaction: Application to bioprosthetic heart valves, Computer Methods in Applied Mechanics and Engineering 284 (2015) 1005–1053.

[18] C. Habchi, S. Russeil, D. Bougeard, J.-L. Harion, T. Lemenand,A. Ghanem, D. Della Valle, H. Peerhossaini, Partitioned solver for strongly coupled fluid–structure interaction, Computers & Fluids 71 (2013) 306–319.

[19] W. Kim, I. Lee, H. Choi, A weak-coupling immersed boundary method for fluid–structure interaction with low density ratio of solid to fluid, Journal of Computational Physics 359 (2018) 296–311.

[20] I. Borazjani, L. Ge, F. Sotiropoulos, Curvilinear immersed boundary method for simulating fluid structure interaction with complex 3D rigid bodies, Journal of Computational Physics 227 (16) (2008) 7587–7620.

[21] H. G. Matthies, J. Steindorf, Partitioned strong coupling algorithms for fluid–structure interaction, Computers & Structures 81 (8-11) (2003) 805–812.

[22] C. W. Hirt, A. A. Amsden, J. Cook, An Arbitrary Lagrangian-Eulerian computing method for all flow speeds, Journal of Computational Physics 14 (3) (1974) 227–253.

[23] M. S. Gadala, M. Movahhedy, J. Wang, On the mesh motion for ALE modeling of metal forming processes, Finite Elements in Analysis and Design 38 (5) (2002) 435–459.

[24] A. Shamanskiy, B. Simeon, Mesh moving techniques in fluid-structure interaction: robustness, accumulated distortion and computational efficiency, Computational Mechanics 67 (2) (2021) 583–600.

[25] A. Leuprecht, S. Kozerke, P. Boesiger, K. Perktold, Blood flow in the human ascending aorta: a combined MRI and CFD study, Journal of Engineering Mathematics 47 (3) (2003) 387–404.

[26] P. Le Tallec, J. Mouro, Fluid structure interaction with large structural displacements, Computer Methods in Applied Mechanics and Engineering 190 (24-25) (2001) 3039–3067.

[27] M. Alishahi, M. Alishahi, H. Emdad, Numerical simulation of blood flow in a flexible stenosed abdominal real aorta, Scientia Iranica 18 (6) (2011) 1297–1305.

[28] A. Qiao, W. Yin, B. Chu, Numerical simulation of fluid–structure interaction in bypassed debakey III aortic dissection, Computer Methods in Biomechanics and Biomedical Engineering 18 (11) (2015) 1173–1180.

[29] K. Bäumler, V. Vedula, A. M. Sailer, J. Seo, P. Chiu, G. Mistelbauer,F. P. Chan, M. P. Fischbein, A. L. Marsden, D. Fleischmann, Fluid– structure interaction simulations of patient-specific aortic dissection, Biomechanics and Modeling in Mechanobiology 19 (5) (2020) 1607–1628.

[30] T. Kim, P. A. J. van Bakel, N. Nama, N. Burris, H. J. Patel, D. M. Williams, C. A. Figueroa, A computational study of dynamic obstruction in type B aortic dissection, Journal of Biomechanical Engineering 145 (3) (2023) 031008.

[31] C. S. Peskin, The immersed boundary method, Acta Numerica 11 (2002) 479–517.

[32] Y. Kim, C. S. Peskin, Penalty immersed boundary method for an elastic boundary with mass, Physics of Fluids 19 (5) (2007).

[33] B. E. Griffith, C. S. Peskin, On the order of accuracy of the immersed boundary method: Higher order convergence rates for sufficiently smooth problems, Journal of Computational Physics 208 (1) (2005) 75– 105.

[34] R. Mittal, H. Dong, M. Bozkurttas, F. Najjar, A. Vargas,A. Von Loebbecke, A versatile sharp interface immersed boundary method for incompressible flows with complex boundaries, Journal of Computational Physics 227 (10) (2008) 4825–4852.

[35] Y.-H. Tseng, J. H. Ferziger, A ghost-cell immersed boundary method for flow in complex geometry, Journal of Computational Physics 192 (2) (2003) 593–623.

[36] M. Uhlmann, An immersed boundary method with direct forcing for the simulation of particulate flows, Journal of Computational Physics 209 (2) (2005) 448–476.

[37] L. Ge, F. Sotiropoulos, A numerical method for solving the 3D unsteady incompressible Navier–Stokes equations in curvilinear domains with complex immersed boundaries, Journal of Computational Physics 225 (2) (2007) 1782–1809.

[38] M. De Tullio, A. Cristallo, E. Balaras, R. Verzicco, Direct numerical simulation of the pulsatile flow through an aortic bileaflet mechanical heart valve, Journal of Fluid Mechanics 622 (2009) 259–290.

[39] B. E. Griffith, X. Luo, D. M. McQueen, C. S. Peskin, Simulating the fluid dynamics of natural and prosthetic heart valves using the immersed boundary method, International Journal of Applied Mechanics 1 (01) (2009) 137–177.

[40] M. Astorino, J.-F. Gerbeau, O. Pantz, K.-F. Traoré, Fluid–structure interaction and multi-body contact: application to aortic valves, Computer Methods in Applied Mechanics and Engineering 198 (45-46) (2009) 3603–3612.

[41] I. Zreid, R. Behnke, M. Kaliske, ALE formulation for thermomechanical inelastic material models applied to tire forming and curing simulations, Computational Mechanics 67 (6) (2021) 1543–1557.

[42] G. L. Nicolson, The Fluid—Mosaic model of membrane structure: Still relevant to understanding the structure, function and dynamics of biological membranes after more than 40 years, Biochimica et Biophysica Acta (BBA)-Biomembranes 1838 (6) (2014) 1451–1466.

[43] A. Bonito, R. H. Nochetto, M. S. Pauletti, Parametric FEM for geometric biomembranes, Journal of Computational Physics 229 (9) (2010) 3171–3188.

[44] T. Kim, R. Ortigosa, N. Nama, M. Aguirre, A. J. Gil, J. D. Humphrey, C. A. Figueroa, A systematic comparison of membrane, shell, and 3d solid formulations for nonlinear vascular biomechanics, Journal of the Mechanical Behavior of Biomedical Materials (2026) 107423.

[45] A. Gilmanov, T. B. Le, F. Sotiropoulos, A numerical approach for simulating fluid structure interaction of flexible thin shells undergoing arbitrarily large deformations in complex domains, Journal of Computational Physics 300 (2015) 814–843.

[46] N. Nama, M. Aguirre, J. D. Humphrey, C. A. Figueroa, A nonlinear rotation-free shell formulation with prestressing for vascular biomechanics, Scientific Reports 10 (1) (2020) 17528.

[47] A. Nitti, J. Kiendl, A. Reali, M. D. de Tullio, An immersed-boundary/isogeometric method for fluid–structure interaction involving thin shells, Computer Methods in Applied Mechanics and Engineering 364 (2020) 112977.

[48] J. Boustani, M. F. Barad, C. C. Kiris, C. Brehm, An immersed boundary fluid–structure interaction method for thin, highly compliant shell structures, Journal of Computational Physics 438 (2021) 110369.

[49] C. A. Taylor, T. J. Hughes, C. K. Zarins, Finite element modeling of blood flow in arteries, Computer Methods in Applied Mechanics and Engineering 158 (1-2) (1998) 155–196.

[50] C. A. Taylor, C. Figueroa, Patient-specific modeling of cardiovascular mechanics, Annual review of Biomedical Engineering 11 (1) (2009) 109– 134.

[51] L. Zhang, A. Gerstenberger, X. Wang, W. K. Liu, Immersed finite element method, Computer Methods in Applied Mechanics and Engineering 193 (21-22) (2004) 2051–2067.

[52] W. K. Liu, Y. Liu, D. Farrell, L. Zhang, X. S. Wang, Y. Fukui, N. Patankar, Y. Zhang, C. Bajaj, J. Lee, et al., Immersed finite element method and its applications to biological systems, Computer Methods in Applied Mechanics and Engineering 195 (13-16) (2006) 1722–1749.

[53] C. J. Arthurs, R. Khlebnikov, A. Melville, M. Marčan, A. Gomez, D. Dillon-Murphy, F. Cuomo, M. Silva Vieira, J. Schollenberger, S. R. Lynch, et al., CRIMSON: An open-source software framework for cardiovascular integrated modelling and simulation, PLoS Computational Biology 17 (5) (2021) e1008881.

[54] C. H. Whiting, K. E. Jansen, A stabilized finite element method for the incompressible Navier–Stokes equations using a hierarchical basis, International Journal for Numerical Methods in Fluids 35 (1) (2001) 93–116.

[55] L. P. Franca, S. L. Frey, Stabilized finite element methods: Ii. the incompressible navier-stokes equations, Computer Methods in Applied Mechanics and Engineering 99 (2-3) (1992) 209–233.

[56] K. E. Jansen, C. H. Whiting, G. M. Hulbert, A generalized-α method for integrating the filtered navier–stokes equations with a stabilized finite element method, Computer methods in applied mechanics and engineering 190 (3-4) (2000) 305–319.

[57] Simmetrix Inc., Simmetrix simulation modeling suite, https://www.simmetrix.com/, Accessed: 2026-05-05 (2026).

[58] T. Belytschko, W. K. Liu, B. Moran, K. Elkhodary, Nonlinear finite elements for continua and structures, John Wiley & Sons, 2014.

[59] M. Bischoff, K.-U. Bletzinger, W. Wall, E. Ramm, Models and finite elements for thin-walled structures, Encyclopedia of Computational Mechanics (2004).

[60] Z. Liu, W. Hong, Z. Suo, S. Swaddiwudhipong, Y. Zhang, Modeling and simulation of buckling of polymeric membrane thin film gel, Computational Materials Science 49 (1) (2010) S60–S64.

[61] E. Lac, D. Barthes-Biesel, N. Pelekasis, J. Tsamopoulos, Spherical capsules in three-dimensional unbounded Stokes flows: effect of the membrane constitutive law and onset of buckling, Journal of Fluid Mechanics 516 (2004) 303–334.

[62] M. Lenz, D. J. Crow, J.-F. Joanny, Membrane buckling induced by curved filaments, Physical review letters 103 (3) (2009) 038101.

[63] J. D. Humphrey, Cardiovascular solid mechanics: cells, tissues, and organs, Springer Science & Business Media, 2013.

[64] Y.-c. Fung, Biomechanics: mechanical properties of living tissues, Springer Science & Business Media, 2013.

[65] C. Geuzaine, J.-F. Remacle, Gmsh: A 3-D finite element mesh generator with built-in pre-and post-processing facilities, International Journal for Numerical Methods in Engineering 79 (11) (2009) 1309–1331.

[66] J. A. Cottrell, T. J. Hughes, Y. Bazilevs, Isogeometric analysis: toward integration of CAD and FEA, John Wiley & Sons, 2009.

[67] J. Chung, G. Hulbert, A time integration algorithm for structural dynamics with improved numerical dissipation: the generalized-α method, Journal of Applied Mechanics 60 (2) (1993) 371–375.

[68] M. Vanella, E. Balaras, A moving-least-squares reconstruction for embedded-boundary formulations, Journal of Computational Physics 228 (18) (2009) 6617–6628.

[69] J. Capecelatro, O. Desjardins, An Euler–Lagrange strategy for simulating particle-laden flows, Journal of Computational Physics 238 (2013) 1–31.

[70] R. W. Hockney, J. W. Eastwood, Computer simulation using particles, crc Press, 2021.

[71] M. D. de Tullio, G. Pascazio, A moving-least-squares immersed boundary method for simulating the fluid–structure interaction of elastic bodies with arbitrary thickness, Journal of Computational Physics 325 (2016) 201–225.

[72] P. Moin, J. Kim, Numerical investigation of turbulent channel flow, Journal of Fluid Mechanics 118 (1982) 341–377.

[73] J. Yang, E. Balaras, An embedded-boundary formulation for large-eddy simulation of turbulent flows interacting with moving boundaries, Journal of Computational Physics 215 (1) (2006) 12–40.

[74] E. P. Mücke, I. Saias, B. Zhu, Fast randomized point location without preprocessing in two-and three-dimensional Delaunay triangulations, in: Proceedings of the twelfth annual symposium on Computational geometry, 1996, pp. 274–283.

[75] M. A. Fernández, Coupling schemes for incompressible fluid-structure interaction: implicit, semi-implicit and explicit, SeMA Journal 55 (1) (2011) 59–108.

[76] C. Farhat, K. G. Van der Zee, P. Geuzaine, Provably second-order time-accurate loosely-coupled solution algorithms for transient nonlinear computational aeroelasticity, Computer Methods in Applied Mechanics and Engineering 195 (17-18) (2006) 1973–2001.

[77] R. D. Rausch, J. T. Batina, H. T. Yang, Euler flutter analysis of airfoils using unstructured dynamic meshes, Journal of Aircraft 27 (5) (1990) 436–443.

[78] C. Farhat, M. Lesoinne, Two efficient staggered algorithms for the serial and parallel solution of three-dimensional nonlinear transient aeroelastic problems, Computer Methods in Applied Mechanics and Engineering 182 (3-4) (2000) 499–515.

[79] S. Rugonyi, K.-J. Bathe, On finite element analysis of fluid flows fully coupled with structural interactions, CMES-Computer Modeling in Engineering and Sciences 2 (2) (2001) 195–212.

[80] W. A. Wall, D. P. Mok, E. Ramm, Partitioned analysis approach of the transient coupled response of viscous fluids and flexible structures, in: Solids, structures and coupled problems in engineering, proceedings of the European conference on computational mechanics ECCM, Vol. 99, 1999, p. 182.

[81] C. Förster, W. A. Wall, E. Ramm, Artificial added mass instabilities in sequential staggered coupling of nonlinear structures and incompressible viscous flows, Computer Methods in Applied Mechanics and Engineering 196 (7) (2007) 1278–1293.

[82] P. Causin, J.-F. Gerbeau, F. Nobile, Added-mass effect in the design of partitioned algorithms for fluid–structure problems, Computer Methods in Applied Mechanics and Engineering 194 (42-44) (2005) 4506–4527.

[83] U. Küttler, M. Gee, C. Förster, A. Comerford, W. Wall, Coupling strategies for biomedical fluid–structure interaction problems, International Journal for Numerical Methods in Biomedical Engineering 26 (3-4) (2010) 305–321.

[84] B. M. Irons, R. C. Tuck, A version of the Aitken accelerator for computer iteration, International Journal for Numerical Methods in Engineering 1 (3) (1969) 275–277.

[85] U. Küttler, W. A. Wall, Fixed-point fluid–structure interaction solvers with dynamic relaxation, Computational Mechanics 43 (1) (2008) 61–72.

[86] T. He, On a partitioned strong coupling algorithm for modeling fluid– structure interaction, International Journal of Applied Mechanics 7 (02) (2015) 1550021.

[87] A. Shenoy, C. Kleinstreuer, Flow over a thin circular disk at low to moderate reynolds numbers, Journal of Fluid Mechanics 605 (2008) 253– 262.

[88] F.-B. Tian, H. Dai, H. Luo, J. F. Doyle, B. Rousseau, Fluid–structure interaction involving large deformations: 3D simulations and applications to biological systems, Journal of Computational Physics 258 (2014) 451– 469.

[89] F. W. Roos, W. W. Willmarth, Some experimental results on sphere and disk drag, AIAA journal 9 (2) (1971) 285–291.

[90] W.-X. Huang, H. J. Sung, Three-dimensional simulation of a flapping flag in a uniform flow, Journal of Fluid Mechanics 653 (2010) 301–336.

[91] W. Nichols, Mcdonald’s blood flow in arteries, Theoretical, Experimental and Clinical Principles 67 (2005) 321–337.

[92] S. Roccabianca, C. Figueroa, G. Tellides, J. D. Humphrey, Quantification of regional differences in aortic stiffness in the aging human, Journal of the Mechanical Behavior of Biomedical Materials 29 (2014) 618–634.

[93] D. Dillon-Murphy, A. Noorani, D. Nordsletten, C. A. Figueroa, Multimodality image-based computational analysis of haemodynamics in aortic dissection, Biomechanics and Modeling in Mechanobiology 15 (4) (2016) 857–876.

